# Genome-wide biases in the rate and molecular spectrum of spontaneous mutations in *Vibrio cholerae* and *Vibrio fischeri*

**DOI:** 10.1101/066829

**Authors:** Marcus M. Dillon, Way Sung, Robert Sebra, Michael Lynch, Vaughn S. Cooper

**Affiliations:** Microbiology Graduate Program, University of New Hampshire, Durham, NH, USA; Department of Bioinformatics and Genomics, University of North Carolina Charlotte, Charlotte, NC, USA; Department of Biology, Indiana University, Bloomington, IN, USA; Department of Microbiology and Molecular Genetics, University of Pittsburgh School of Medicine, Pittsburgh, PA, USA; Department of Genetics and Genomic Sciences and Icahn Institute for Genomics and Multiscale Biology, Icahn School of Medicine at Mount Sinai, New York, NY, USA

**Keywords:** *Vibrio cholerae*, *Vibrio fischeri*, mutation accumulation, mutation rate, mutation spectra

## Abstract

The vast diversity in nucleotide composition and architecture among bacterial genomes may be partly explained by inherent biases in the rates and spectra of spontaneous mutations. Bacterial genomes with multiple chromosomes are relatively unusual but some are relevant to human health, none more so than the causative agent of cholera, *Vibrio cholerae*. Here, we present the genome-wide mutation spectra in wild-type and mismatch repair (MMR) defective backgrounds of two *Vibrio* species, the low-GC% squid symbiont *V. fischeri* and the pathogen *V. cholerae*, collected under conditions that greatly minimize the efficiency of natural selection. In apparent contrast to their high diversity in nature, both wild-type *V. fischeri* and *V. cholerae* have among the lowest rates for base-substitution mutations (bpsms) and insertion-deletion mutations (indels) that have been measured, below 10^−3^/genome/generation. *V. fischeri* and *V. cholerae* have distinct mutation spectra, but both are AT-biased and produce a surprising number of multi-nucleotide indels. Furthermore, the loss of a functional MMR system caused the mutation spectra of these species to converge, implying that the MMR system itself contributes to species-specific mutation patterns. Bpsm and indel rates varied among genome regions, but do not explain the more rapid evolutionary rates of genes on chromosome 2, which likely result from weaker purifying selection. More generally, the very low mutation rates of *Vibrio* species correlate inversely with their immense population sizes and suggest that selection may not only have maximized replication fidelity but also optimized other polygenic traits relative to the constraints of genetic drift.

## INTRODUCTION

In 2000, the complete genome sequence of *Vibrio cholerae*, the first bacterial genome with multiple chromosomes to be completed, was published in *Nature* to acclaim (Heidelberg et al. 2000). Deemed a “treasure trove” for microbiology and genomics researchers (Waldor and RayChaudhuri 2000), the *V. cholerae* genome revealed several intriguing asymmetries between the chromosomes. The larger chromosome 1 (chr1) is approximately 3 Mb in size and contains most essential genes and the pathogenicity elements required to cause the disease cholera, whereas the smaller chromosome 2 (chr2) is approximately 1 Mb and contains few essential genes but many undefined genes acquired by horizontal transfer (Heidelberg et al. 2000; Cooper et al. 2010). The plasmid-like origin of replication on chr2 implies that it originated as a megaplasmid that became entrapped by the translocation of essential genes, and the greater variation in gene content among strains on chr2 has led many to speculate that the small chromosome somehow confers an evolutionary advantage in varied environments, perhaps by being enriched for conditionally useful traits (Schoolnik and Yildiz 2000). Indeed, this two-chromosome structure is found throughout the *Vibrionaceae* family, suggesting an uncertain evolutionary advantage in their aquatic habitats.

Subsequent studies of the evolution of *Vibrio* genomes have shown that not only is chr2 more variable in its content, its conserved genes tend to evolve more rapidly (Cooper et al. 2010). These elevated evolutionary rates likely stem from a gene dosage bias between chr1 and chr2 that emerges from their replication timing. Specifically, replication of chr2 occurs later than chr1, leading to synchronous termination of replication of both chromosomes, even though chr1 is larger (Egan and Waldor 2003; Duigou et al. 2006; Rasmussen et al. 2007). The earlier replication of two-thirds of chr1 generates a gene dosage bias that can be greatly compounded during rapid growth in which multiple replication forks are active (Stokke et al. 2011). Consequently, genes replicated early tend to be expressed more, experience greater purifying selection, and hence evolve more slowly (Cooper et al. 2010; Morrow and Cooper 2012). Effects of translocations between chromosomes also support this model: genes moving between chromosomes tend to exhibit altered expression rates that match their new neighbors (Morrow and Cooper 2012), and orthologs found on secondary chromosomes are predisposed to more rapid evolution even when found in related genomes with a single chromosome (Cooper et al. 2010).

While these comparative-genomic studies suggest that replication timing and its effects on gene dosage may influence the strength of purifying selection (and hence the fate of genetic variation), evidence also exists that replication timing positively correlates, albeit weakly, with mutation rates (the origin of variation) in all domains of life (Stamatoyannopoulos et al. 2009; Chen et al. 2010; Agier and Fischer 2012; Martincorena et al. 2012). Mutation rates may be non-uniform because of greater rates of damage, asymmetric nucleotide pools, structural differences affecting polymerase fidelity, biases in gene expression or replication, or variation in the effectiveness of DNA repair (Sharp et al. 1989; Zhang and Mathews 1995; Mira and Ochman 2002; Cooper et al. 2010; Lee et al. 2012). If late-replicated genes experience both weaker purifying selection and greater mutation rates relative to early-replicated genes, selection could therefore act on genome organization to influence both the origin and fate of genetic variation.

The goal of this study was to examine how mutation rates and spectra vary within and among chromosomes based on existing models of replication timing in two species of *Vibrio, V. cholerae* and *V. fischeri*. Both species are globally significant and are often compared to one another because of their marine habitats and similar genome structures, but contrasting roles in pathogenesis and symbiosis (Goldberg and Murphy 1983; Thompson et al. 2004; Ruby et al. 2005). We also evaluated how the mismatch repair (MMR) system influenced the rates and spectra of spontaneous mutations in both organisms by removing the *mutS* gene from each genome. Our approach used the well-documented method of mutation accumulation (MA) paired with whole-genome sequencing (WGS) (Behringer and Hall 2016). Using the MA-WGS method, a single clonal ancestor is used to initiate many replicate lineages that are passaged through hundreds of single cell bottlenecks before each being subjected to WGS. The bottlenecking regime minimizes the efficiency of natural selection on mutations, and when the genome sequences are compared, a nearly unbiased picture of the natural mutation rates and spectra of the ancestor is revealed. A growing body of MA-WGS studies in microbes have revealed some general properties of spontaneous mutation (Lynch et al. 2008; Denver et al. 2009; Ossowski et al. 2010; Lee et al. 2012; Sung et al. 2012; Schrider et al. 2013; Heilbron et al. 2014; Long et al. 2014; Zhu et al. 2014; Dillon et al. 2015; Long et al. 2015; Sung et al. 2015; Dettman et al. 2016), but unique properties of the spontaneous mutation rates and spectra in many taxa have revealed the value of conducting detailed MA-WGS studies in a more comprehensive collection of species, particularly those with multiple chromosomes.

Here, we performed four detailed MA-WGS experiments on both wild-type and MMR-deficient strains of *V. fischeri* and *V. cholerae*. We identified a total of 439 wild-type mutations and 5990 *ΔmutS* mutations distributed across both chromosomes of *V. fischeri* and *V. cholerae*, producing the first genome-wide perspective of the bpsm and indel biases in wild-type and MMR-deficient strains of these two significant bacterial species. Mutation rates were non-uniform among genome regions and varied in patterns that appear to be associated with replication timing. Both genomes also exhibited distinct base-substitution mutation spectra that correlated with their different %GC content and also experienced a surprising number of multi-nucleotide indel mutations. Remarkably, the loss of MMR caused both mutation spectra to converge towards similar patterns dominated by transition mutations and small indels, which suggests that variation in the MMR system may be partly responsible for producing differences in genome composition.

## RESULTS

The two ancestral genomes for this study share the two-chromosome architecture common to *Vibrio* but differ in other important aspects. The genome of *V. fischeri* strain ES114, which was completely sequenced previously, harbors two chromosomes of 2.90 Mb and 1.33 Mb that differ slightly in their %GC content (38.96% for chr1 and 37.02% for chr2). In addition, *V. fischeri* ES114 has a 45.85 Kb plasmid with a %GC content of 38.44% (Ruby et al. 2005). The genome of *V. cholerae* strain 2740–80, which was previously deposited as a draft assembly, was closed and polished for this study using a combination of PacBio sequencing, Illumina sequencing, and consensus assemblies (see Methods). This *V. cholerae* genome includes a 2.99 Mb primary chromosome (chr1) and a 1.10 Mb secondary chromosome (chr2), with %GC contents of 47.85% and 46.83%, respectively. Four MA experiments were conducted with these strains using daily single-cell bottlenecks produced by isolation-streaking on agar plates, which limits the efficiency of natural selection to purge deleterious and enrich beneficial mutations. For the two wild-type (wt) experiments, *V. fischeri* ES114 (*Vf*-wt) and *V. cholerae* 2740–80 (*Vc*-wt) colonies were used to found 75 MA lineages, each of which was propagated for 217 days. For the two mutator (mut) experiments, *V. fischeri* ES114 (*Vf*-mut) and *V. cholerae* 2740–80 (*Vc*-mut) strains lacking a *mutS* gene were used to found 48 MA lineages, each of which was propagated for 43 days. The properties of each of these experiments, the observed mutation spectra, and the lack of genetic parallelism suggest that few of these mutations were subject to the biases of natural selection (Supplemental Text). Summary parameters of each MA experiment, including the numbers of mutations identified in all final isolates, generations of growth, and overall mutation rates are summarized in Table 1. Owing to the fitness cost of increasing mutational load, generations of growth per day declined over the course of each MA experiment, particularly in the mutator lineages (Figure S1).

**Table 1.** Parameters and observed mutations in the four mutation accumulation experiments.

| MA Lines | Sequenced Lines | Gen. per line | Gen. total | No. of bpsm | No. of indels | Bpsm rate per nucleotide<sup>a</sup> | Bpsm rate per genome<sup>b</sup> | Indel rate per nucleotide<sup>a</sup> | Indel rate per genome<sup>b</sup> |
| --- | --- | --- | --- | --- | --- | --- | --- | --- | --- |
| Vf_wt | 48 | 5187 | 248976 | 219 | 60 | $2.07 \times 10^{-10}$ | $8.85 \times 10^{-4}$ | $5.68 \times 10^{-11}$ | $2.43 \times 10^{-4}$ |
| Vc_wt | 49 | 6453 | 316197 | 138 | 22 | $1.07 \times 10^{-10}$ | $4.38 \times 10^{-4}$ | $1.71 \times 10^{-11}$ | $6.98 \times 10^{-5}$ |
| Vf_mut | 19 | 810 | 15390 | 4313 | 382 | $6.57 \times 10^{-8}$ | $2.81 \times 10^{-1}$ | $5.82 \times 10^{-9}$ | $2.49 \times 10^{-2}$ |
| Vc_mut | 22 | 1254 | 27588 | 1022 | 273 | $9.09 \times 10^{-9}$ | $3.72 \times 10^{-2}$ | $2.43 \times 10^{-9}$ | $9.93 \times 10^{-3}$ |

<sup>a</sup>Bpsm and indel mutation rates/nucleotide/generation are calculated as the number of observed mutations, divided by the product of sites analyzed and number of generations per lineage. The above estimates represent the average rate across all sequenced lineages.
<sup>b</sup>Bpsm and indel mutation rates/genome/generation are calculated by multiplying the mutation rate/nucleotide/generation in each lineage by the genome size. The above estimates represent the average rate across all sequenced lineages.

### Wild-type base-substitution mutation rates and spectra

The wild-type base-substitution mutation rates for these *Vibrio* species are among the lowest per-generation rates that have been observed in any organism (Figure S2). The bpsm rate for *V. fischeri* is 2.07 (0.207) • 10^−10^ /bp/generation (SEM), which is approximately twice the rate of bpsm in *V. cholerae*, 1.07 (0.094) • 10^−10^ /bp/generation (SEM). These per-site bpsm rates correspond to genome-wide bpsm rates of 0.0009 /genome/generation for *V. fischeri* and 0.0004 /genome/generation for *V. cholerae*. (Table 1). Bpsm rates differ somewhat between the chromosomes of *V. fischeri*, with chr2 experiencing significantly greater rates than chr1 (χ^2^ = 4.799, d.f. = 1, p = 0.028), but no difference was found between the chromosomes of *V. cholerae* (χ^2^ = 0.556, d.f. = 1, p = 0.456) (Figure 1A). The increased bpsm rate on chr2 of *V. fischeri* cannot be explained by the relative nucleotide contents of the chromosomes (%GC: chr1, 39.0%; chr2, 37.0%). In fact, when we account for AT-biased mutation in *V. fischeri*, the difference between the expected and observed bpsms between the two chromosomes increased (χ^2^ = 5.747, d.f. = 1, p = 0.017). Correcting for AT-biased mutation in *V. cholerae* (%GC: chr1, 47.9%; chr2, 46.8%) revealed no significant difference between the bpsm rates of chr1 and chr2 (χ^2^ = 0.123, d.f. = 1, p = 0.726).

**Figure 1.**
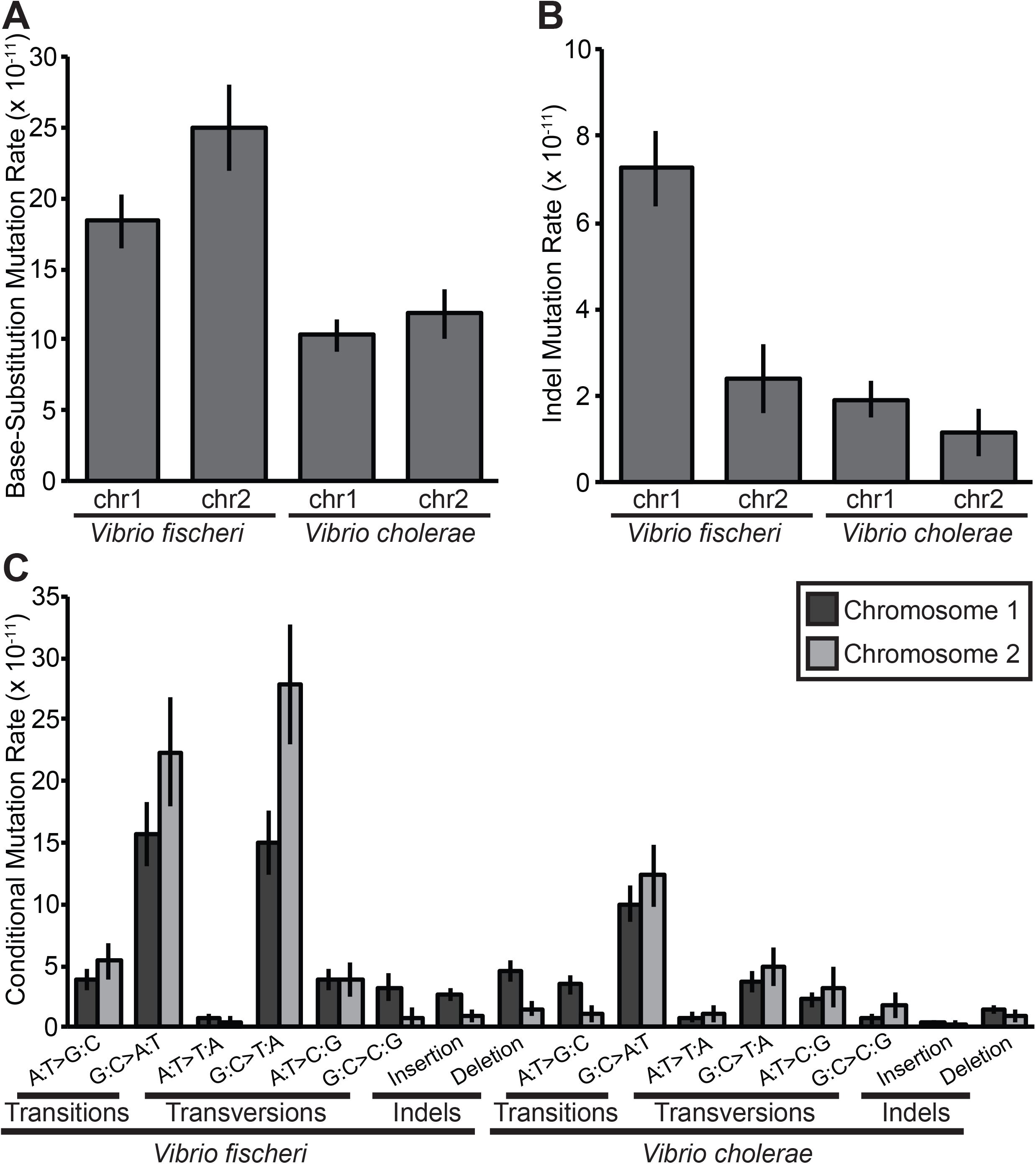
Wild-type base-substitution (bpsm) and insertion-deletion (indel) mutation rates and spectra for the two chromosomes of *Vibrio fischeri* and *Vibrio cholerae*; error bars indicate one standard error of the mean. (A and B) Overall bpsm and indel mutation rates per base-pair per generation. (C) Conditional bpsm and indel rates per conditional base-pair per generation, estimated by dividing the number of observed mutations by the product of the analyzed sites capable of producing a given mutation and the number of generations of mutation accumulation in each lineage.

Base-substitution mutation spectra in both *V. fischeri* and *V. cholerae* were significantly AT-biased *(Vf:* χ^2^ = 108.090, d.f. = 1, p < 0.0001 ; *Vc*: χ^2^ = 28.744, d.f. = 1, p < 0.0001) (Figure 1C). Interestingly, the AT-bias is stronger in *V. fischeri*, which has the lower genome wide %GC-content. However, consistent with previous MA studies of bacteria (Lind and Andersson 2008; Lee et al. 2012; Sung et al. 2012; Dillon et al. 2015; Sung et al. 2015; Dettman et al. 2016), AT-mutation bias alone fails to explain realized genome-wide %GC-contents. For *V. fischeri*, the expected %GC-content under mutation-drift equilibrium is 0.202 ± 0.034 (SEM), 0.182 lower than the genome-wide %GC-content (0.384). This AT-bias is generated by high relative bpsm rates of both G:C > A:T transitions and G:C > T:A transversions, but it is the relative G:C > T:A transversion rate that is especially high in comparison to MA studies in other species. With a rate of 9.19 • 10^−9^/bp/generation, G:C > T:A transversions occur at a higher rate than any other bpsm, generating a transition/transversion ratio (T_S_/T_V_) of 0.851. Similarly, the expected %GC content of *V. cholerae* under mutation-drift equilibrium is 0.281 ± 0.044 (SEM), 0.195 lower than the genome-wide %GC content (0.476). However, in *V. cholerae*, the AT-bias is generated mostly by G:C > A:T transitions rather than G:C > T:A transversions, resulting in a T_S_/T_V_ of 1.59.

To test whether the bpsm spectra on chr1 were significantly different from those on chr2, we used the conditional bpsm rates (corrected for %GC-content) of chr1 to generate expected ratios for each bpsm and compared these with the observed bpsm spectra on chr2 (Figure 1C). Neither the overall bpsm spectra of *V. fischeri* or *V. cholerae* varied significantly between chromosomes (*Vf:* χ^2^ = 7.797, d.f. = 5, p = 0.168; *Vc*: χ^2^ = 6.516, d.f. = 5, p = 0.259). However, the G:C > T:A transversion rate was significantly higher on chr2 of *V. fischeri* (Welch’s two tailed t-test; t = 2.348, df = 71.952, p = 0.022) and the A:T > G:C transition rate was significantly lower on chr2 of *V. cholerae* (Welch’s two tailed t-test; t = −2.155, df = 95.749, p = 0.034) (Figure 1C). Interestingly, late replicating regions of chr1 in *V. fischeri* (the terminal 1,330,333 bp, equal to the size of chr2) also had elevated G:C > T:A transversion rates, and late replicating regions of chr1 in *V. cholerae* (terminal 1,101,931 bp, equal to the size of chr2) had reduced A:T > G:C transition rates relative to early replicating regions, though neither of these intra-chromosomal differences were significant (Welch’s two tailed t-test; *Vf* G:C>T:A T_v_: t = 1.789, df = 81.645, p = 0.077; *Vc* A:T > G:C T_s_: t = −0.720, df = 95.489, p = 0.473). Bpsm rates of other mutation types in late replicating regions of chr1 do not conform to those on chr2 (Table S1), suggesting that not all bpsms are influenced by replication timing.

### Wild-type insertion-deletion mutation rates

Indels occurred at approximately one-fifth the rate of bpsm and occurred predominantly in simple sequence repeats (SSRs). Cumulative indel mutation rates for *V. fischeri* and *V. cholerae* were 5.68 (0.691) • 10^−11^ /bp/generation and 1.71 (0.337) • 10^−11^ /bp/generation (SEM), respectively (Table 1). These rates correspond to genome-wide indel rates of 0.0002 /genome/generation for *V. fischeri* and 0.00007 /genome/generation for *V. cholerae* (Table 1). The indel spectra of both species were biased towards deletions, with deletions occurring at approximately twice the rate of insertions in *V. fischeri* (χ^2^ = 4.267, d.f. = 1, p = 0.039), and three times the rate of insertions in *V. cholerae* (χ^2^ = 6.546, d.f. = 1, p = 0.011) (Figure 1). Indels were also significantly more frequent on chr1 in *V. fischeri* (χ^2^ = 9.066, d.f. = 1, p = 0.003) and slightly more common on chr1 in *V. cholerae* (χ^2^ = 0.123, d.f. = 1, p = 0.726), though not significantly so (Figure 1B).

In contrast with previous reports from bacterial MA-WGS studies, we find many multi-nucleotide indels in both the *Vf*-wt and *Vc*-wt MA experiments. In *Vf*-wt, only 20.00% of indels involved the insertion or deletion of a single nucleotide, while 36.36% of the *Vc*-wt indels involved a single nucleotide. Furthermore, the distribution of the number of indels identified for each length below 10-bps demonstrates that short multi-nucleotide indels were relatively common, particularly in *V. fischeri* (Figure 2).

**Figure 2.**
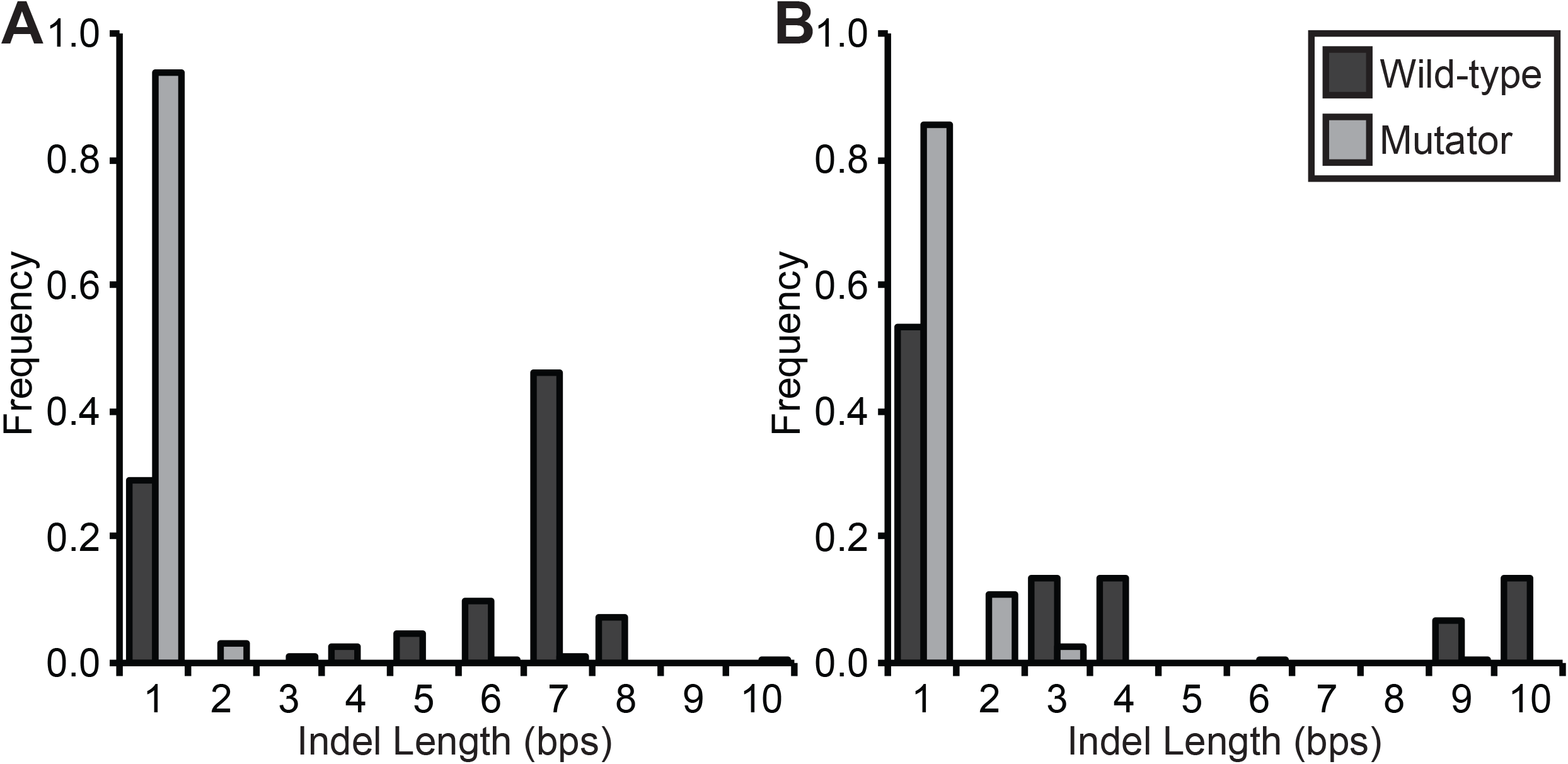
Relative frequency of insertion-deletion mutations (indels) of different lengths observed in the wild-type (wt) and mismatch repair deficient (mut) strains of *Vibrio fischeri* (A) and *Vibrio cholerae* (B). Note that the overall indel rates of *Vf*-mut and *Vc*-mut are 102-fold and 142-fold higher than the wild-type rates, respectively, but it is the relative frequencies of different indel lengths that are shown here. The increase in the number of single-nucleotide indels in mutator lines is significant (*Vf*-mut: χ^2^ = 5.460 • 10^3^, d.f. = 1, p < 0.0001; *Vc*-mut: χ^2^ = 2.567 • 10^3^, d.f. = 1, p < 0.0001).

More strikingly, in *V. fischeri*, the indel rates of SSRs scaled positively with the repeat unit length and the number of repeated units. 41 of the 60 indels (68.33%) observed in *V. fischeri* occurred in SSRs containing three or more repeated units, which is significantly more than we expect based on the frequency of bases in SSRs in the *V. fischeri* genome (χ^2^ = 82.915, d.f. = 1, p < 0.0001). When SSRs are categorized by their repeat unit length (mono-, di-, tri-nucleotide, etc.), SSRs with longer repeat units experience elevated mutation rates (Figure 3). Importantly, these rates vary over orders of magnitude and their observed frequencies differ significantly from the null expectation derived from their target sizes (χ^2^ = 2.120 • 10^5^, d.f. = 8, p < 0.0001). SSR indel rates also scale positively with the number of repeats in the SSR and again differ significantly from the null expectation derived from their genome target (Chi-square test, repeat numbers 3–10; χ^2^ = 5.590 • 10^2^, d.f. = 7, p < 0.0001). Lastly, a few SSRs appear to be especially mutagenic because the same locus mutated independently in multiple lineages (Supplementary Dataset 2). We cannot ascertain whether similar SSR biases exist in *V. cholerae* because only 22 indels were observed and only 8 of those were in SSRs with three or more repeats (36.36%). However, the elevated occurrence of indels in SSRs in *V. cholerae* is consistent with our observations in *V. fischeri* (χ^2^ = 4.547, d.f. = 1, p = 0.033).

**Figure 3.**
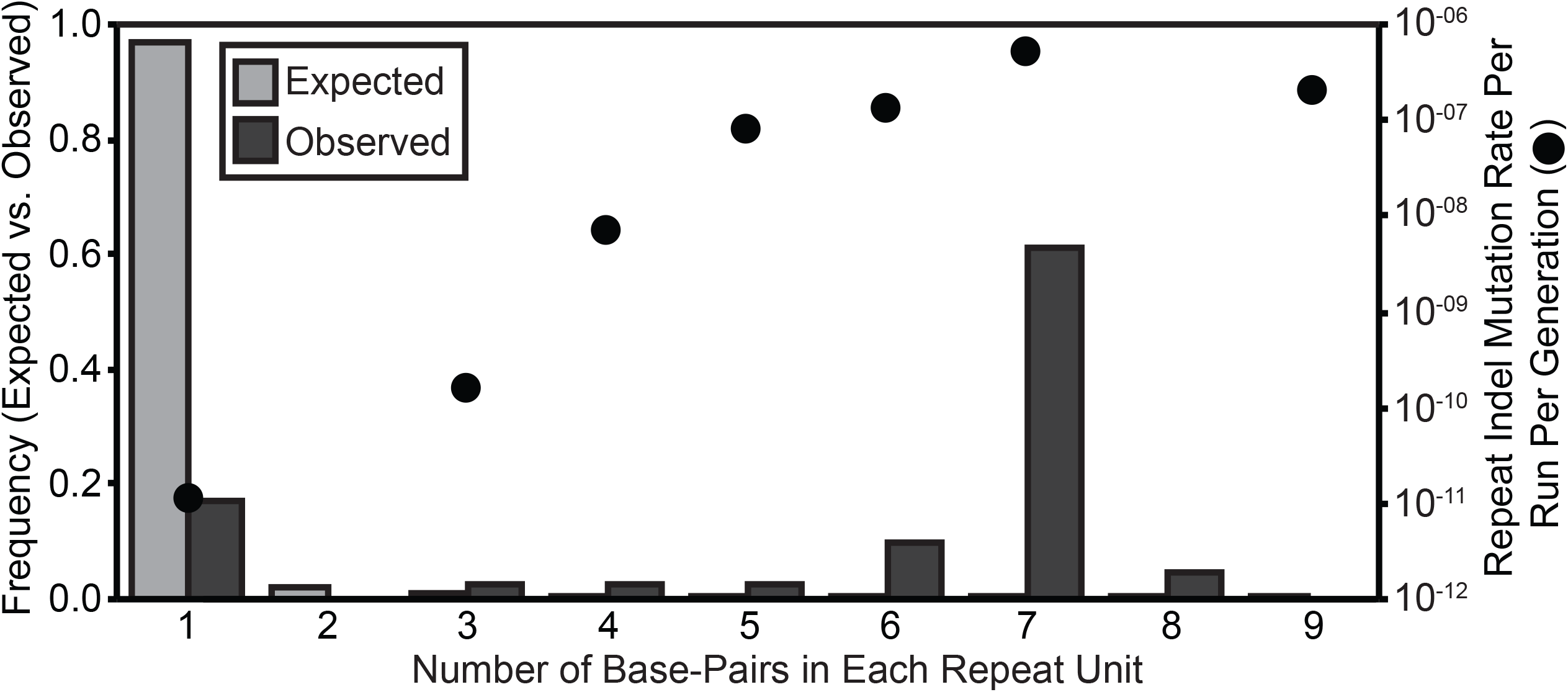
Wild-type insertion-deletion mutation (indel) rates per run per generation and frequencies in simple-sequence repeats (SSRs) containing three or more repeats in *Vibrio fischeri*. SSRs were categorized by the length of the repeated unit and indel rates per run per generation were calculated as the number of observed indels in that SSR category, divided by the product of the occurrence of that SSR type in the genome, the number of generations, and the number of MA lineages analyzed. Expected frequencies were calculated based on the target size of each SSR category in the genome.

### Effects of losing DNA mismatch repair

The deletion of the *mutS* gene results in a faulty MMR system and is expected to increase the rates and alter the spectra of bpsms and indels. In *V. fischeri, ΔmutS* produced a 317-fold increase in the bpsm rate and a 102-fold increase in the indel rate (Table 1). This *ΔmutS* mutation also eliminated differences between chromosomes in both bpsm and indel rates observed in the *Vf*-wt lineages (bpsm: χ^2^ = 0.109, d.f. = 1, p = 0.741; indels: χ^2^ = 2.076, d.f. = 1, p = 0.150). In *V. cholerae*, the *mutS* deletion produced an 85-fold increase in the bpsm rate and a 142-fold increase in the indel rate (Table 1). However, in this case, slightly more bpsm occurred on chr1 in *Vc*-mut lineages than expected (bpsm: χ^2^ = 4.540, d.f. = 1, p = 0.0331; indels: χ^2^ = 0.041, d.f. = 1, p = 0.840).

Losing functional mismatch repair caused the bpsm spectra of both *V. fischeri* and *V. cholerae* to change dramatically and converge (Figure 4). Despite the significantly different bpsm spectra between wild-type species (χ^2^ = 32.788, d.f. = 5, p < 0.0001), the loss of *mutS* caused *Vf*-mut and *Vc*-mut lineages to accumulate a remarkably similar spectrum of bpsms (χ^2^ = 10.178, d.f. = 5, p = 0.070). Specifically, both sets of mutator lines were dominated by A:T > G:C and G:C > T:A transitions, which would yield significantly more %GC-rich genomes of 0.482 ± 0.016 (SEM) in *Vf*-mut and 0.475 ± 0.011 (SEM) in *Vc*-mut at mutation-drift equilibrium. In addition, while *V. fischeri* continues to have a slight deletion bias with defective MMR, insertions actually occur at a slightly higher rate than deletions in the *Vc*-mut lineages, although not significantly so (*Vf*-mut: χ^2^ = 6545, d.f. = 1, p = 0.011; *Vc*-mut: χ^2^ = 0.297, d.f. = 1, p = 0.586). Lastly, as with the wild-type lineages, the bpsm spectra do not vary significantly between chromosomes for either MMR deficient species (*Vf*-mut: χ^2^ = 8.929, d.f. = 1, p = 0.112; *Vc*-mut: χ^2^ = 5.239, d.f. = 1, p = 0.387).

**Figure 4.**
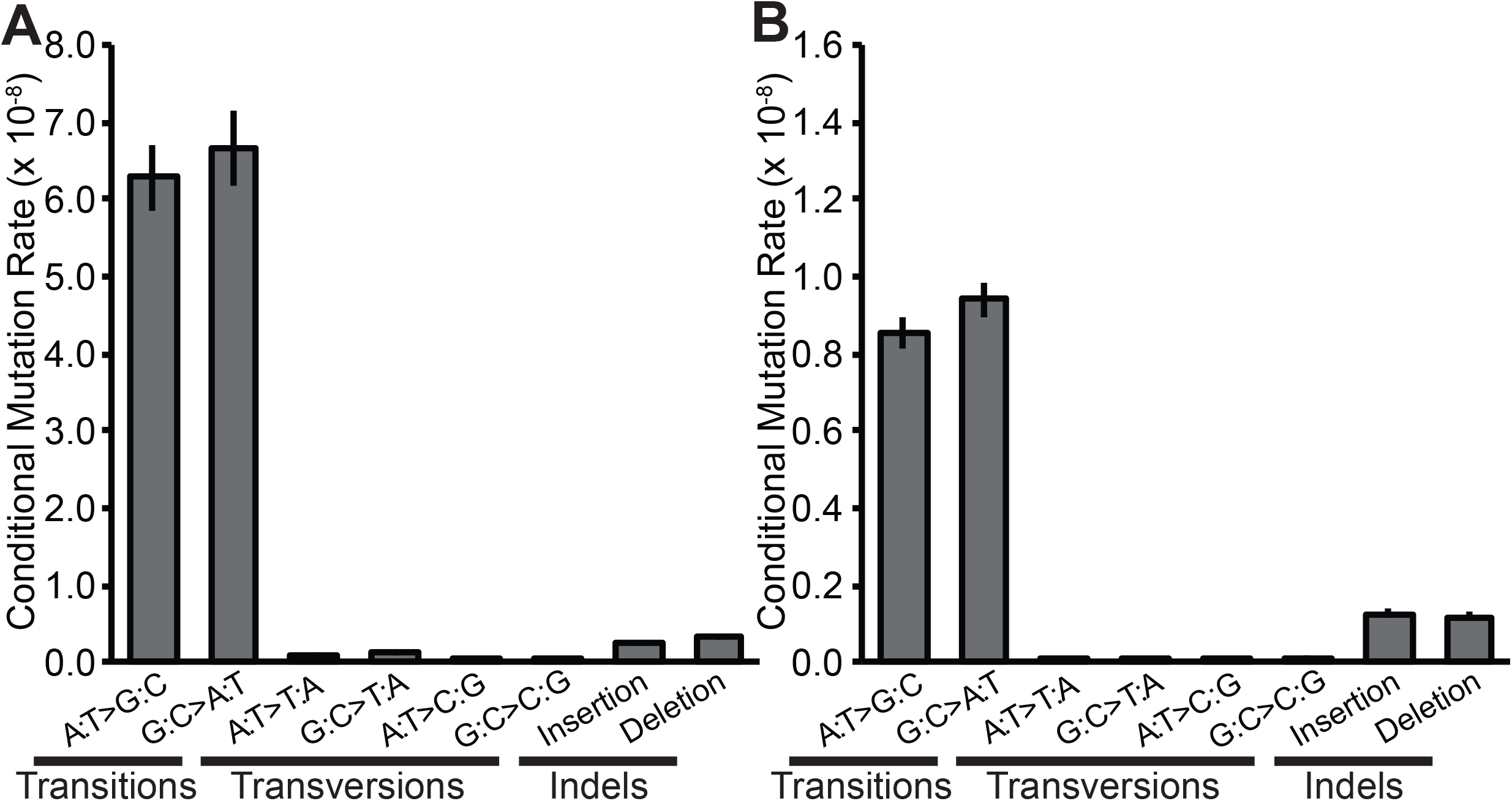
Mismatch repair deficient conditional base-substitution (bpsm) and insertion-deletion (indel) mutation rates per conditional base-pair per generation for *Vibrio fischeri ΔmutS* and *Vibrio cholerae ΔmutS* mutation accumulation lines, estimated by dividing the number of observed mutations by the product of the analyzed sites capable of producing a given mutation and the number of generations of mutation accumulation in each lineage.

While multi-nucleotide indels were relatively common in the wild-type MA lines, the vast majority of indels involved a single nucleotide in both the *Vf*-mut (93.62%) and *Vc*-mut (85.66%) lines (Figure 2). The *mutS* deletion increased the single-nucleotide indel rate 473-fold but the multi-nucleotide indel rate only 13-fold in *V. fischeri*. Similar patterns were seen in *V. cholerae*, where *ΔmutS* increased the single-nucleotide indel rate 334-fold and the multi-nucleotide indel rate only 64-fold. Most of the increase in the multi-nucleotide mutation rate resulted from di-and tri-nucleotide indels that were rarely observed in the wild-type MA experiments (Figure 2). Yet all indels detected in the mutator MA experiments up to 10-bps in length were also significantly over-represented in mutator MA experiments (Table S2), implying that *mutS* and the MMR system in general are involved in the repair of all indel mutations up to 10-bps in length.

Most single-nucleotide indels in *Vf*-mut and *Vc*-mut lines occurred in homopolymeric runs. The single-nucleotide indel mutation rate in both *Vf*-mut and *Vc*-mut correlates positively with the length of the homopolymer (Figure 5) and differs significantly from the null expectation derived from the genome target size (Chi-square test, repeat numbers 3–11; *Vf*-mut: χ^2^ = 3.189 × 10^4^, d.f. = 8, p < 0.0001; *Vc*-mut: χ^2^ = 6.850 × 10^4^, d.f. = 8, p < 0.0001). Because indels in SSRs with longer repeated units were rarely observed in the mutator lineages, we cannot confirm whether the length of the repeated unit in an SSR also correlates positively with its indel rate in either of the mutator lineages.

**Figure 5.**
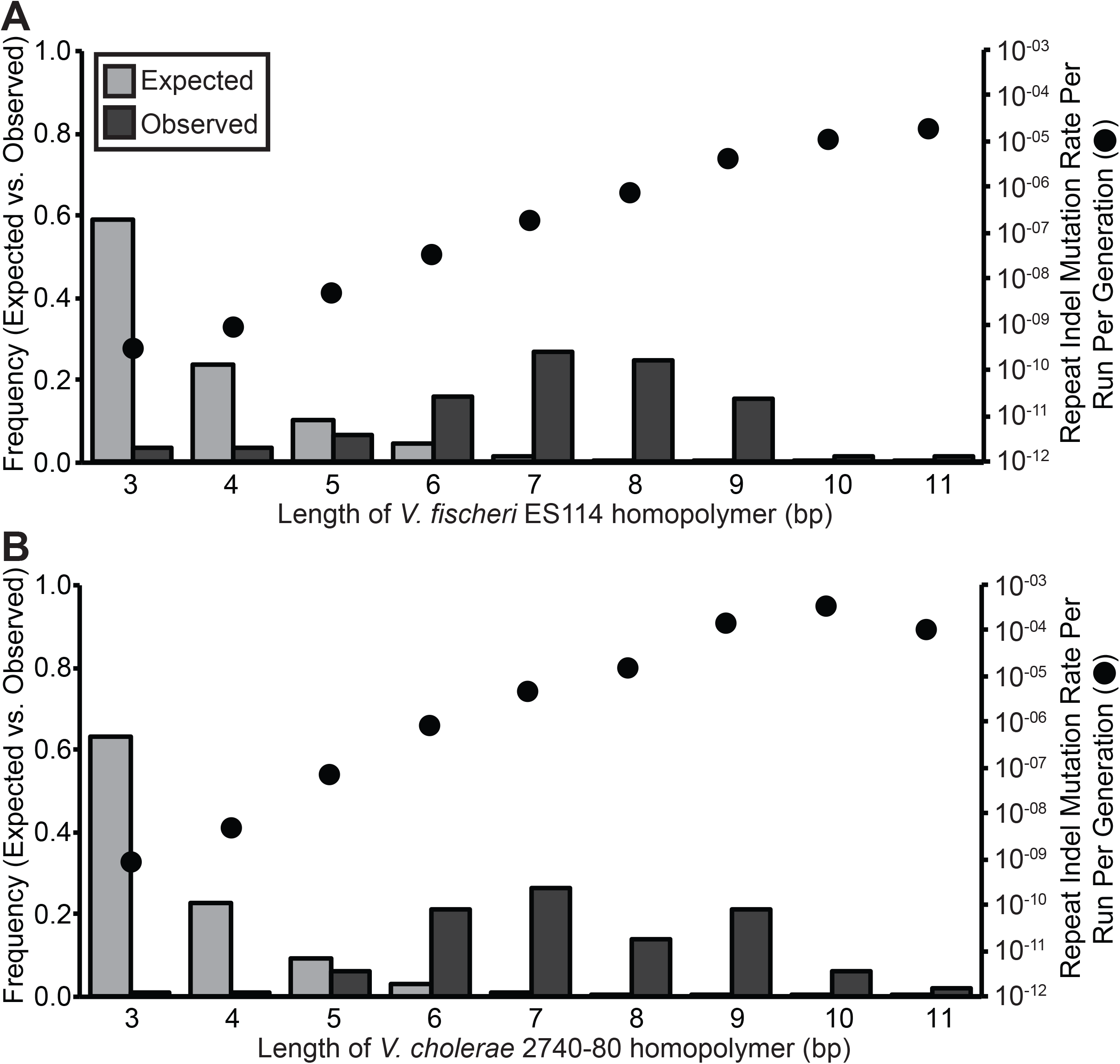
Mismatch repair deficient insertion-deletion mutation (indel) rates per run per generation and frequencies in homopolymer repeats containing three or more repeats for *Vibrio fischeri ΔmutS* (A) and *Vibrio cholerae ΔmutS* (B). Indel rates per run per generation were calculated as the number of observed indels in each homopolymer length category, divided by the product of the occurrence of homopolymers of that length in the genome, the number of generations, and the number of MA lineages analyzed. Expected frequencies were calculated based on the genome target size for each homopolymer length.

### Genomic distribution of spontaneous mutations

To this point we have only discussed inter-chromosomal differences in mutation rates between two autonomously replicating chromosomes. However, the distribution of bpsms and indels in *V. fischeri* and *V. cholerae* may also vary among regions within these circular chromosomes. We analyzed the genome-wide distribution of bpsms by dividing each chromosome into 100 kb intervals extending bi-directionally from the origin of replication. Despite apparent intra-chromosome variation in the bpsm rate of both wild-type ancestors (Figure 6A,B), the observed number of bpsms in 100kb intervals does not differ significantly from random expectations based on the number of analyzed sites (*Vf*-wt Chr1: χ^2^ = 31.254, d.f. = 29, p = 0.354; *Vf*-wt Chr2: χ^2^ = 17.185, d.f. = 15, p = 0.308; *Vc*-wt Chr1: χ^2^ = 29.607, d.f. = 29, p = 0.434; *Vc*-wt Chr2: χ^2^ = 12.825, d.f. = 11, p = 0.305). However, bpsms were not distributed evenly on chr1 in either of the mutator experiments (*Vf*-mut: χ^2^ = 132.970, d.f. = 29, p < 0.0001; *Vc*-mut: χ^2^ = 102.420, d.f. = 29, p < 0.0001), resulting in a mirrored wave-like pattern of bpsm rates on the right and left replichores (Figure 6A,B). Indeed, we find a significant positive relationship between the bpsm rates on the right replichore and concurrently replicated regions on the left replichore in both genomes (Linear regression; *Vf*-mut: F = 10.98, df = 13, p = 0.0060, r^2^ = 0.46; *Vc*-mut: F = 6.76, df = 13, p = 0.0221, r^2^ = 0.34). In contrast to chr1, the observed distribution of bpsms on chr2 of the *V. fischeri* and *V. cholerae* mutator lineages does not differ from a null, constant rate among regions (*Vf*-mut: χ^2^ = 17.459, d.f. = 15, p = 0.292; *Vc*-mut: χ^2^ = 10.984, d.f. = 11, p = 0.445), suggesting that bpsm rates are more consistent across chr2 than they are on chr1 (Figure 6A, B).

**Figure 6.**
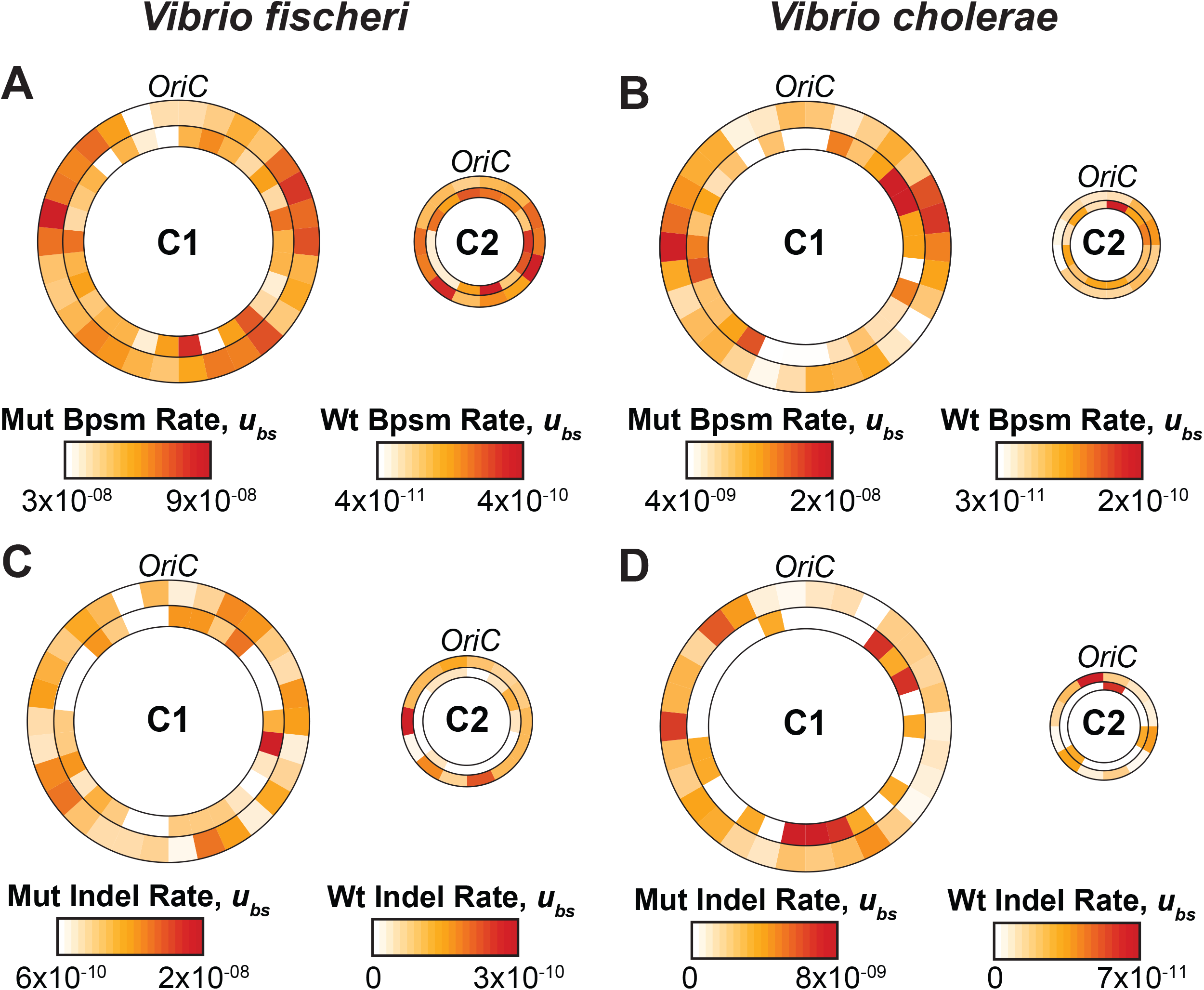
Base-substitution (bpsm) and insertion-deletion (indel) mutation rates per base-pair per generation in 100kb intervals extending bi-directionally from the origin of replication *(OriC)* for all mutation accumulation (MA) experiments in this study. Outer rings on each chromosome represent the mutator MA experiment and inner rings represent the corresponding wild-type MA experiment. A) Bpsm rates of *Vf*-wt and *Vf*-mut; B) Bpsm rates of *Vc*-wt and *Vc*-mut; C) Indel rates of *Vf*-wt and *Vf*-mut; D) Indel rates of *Vc*-wt and *Vc*-mut.

Indels occur predominantly in SSRs, which are not uniformly distributed across the genome, and are thus also expected to vary significantly between genome regions. This variation was seen in all experiments except *Vc*-wt, likely because only 22 indels were detected in this experiment (Figure 6 C,D) (**Vf*-wt*: χ^2^ = 83.079, d.f. = 43, p < 0.0001; *Vc*-wt: χ^2^ = 42.224, d.f. = 41, p = 0.418; *Vf*-mut: χ^2^ = 96.383, d.f. = 43, p < 0.0001; *Vc*-mut: χ^2^ = 157.240, d.f. = 41, p < 0.0001). However, indel rates were not conserved on opposing replichores of either chromosome and were rather dispersed throughout the genomes.

## DISCUSSION

### Causes and effects of high genetic diversity within *Vibrio*

The unique and broadly significant biology of both *V. cholerae* and *V. fischeri* motivated this comprehensive study of their mutational processes in the near-absence of natural selection. *V. cholerae* is one of the most significant human pathogens, infecting 3–5 million people and causing approximately 100,000 deaths annually from side effects of the profuse diarrhea caused by toxigenic strains (Lozano et al Lancet 2012). *V. cholerae* also lives in a broad range of aquatic habitats associated with zooplankton and is often highly abundant, demonstrating that only a small number of strains are capable of causing cholera. This ecological diversity is borne out in the genetic diversity seen in sequenced genomes, namely a high nucleotide diversity at silent sites (πs = 0.110) (Sung et al. 2016). This nucleotide diversity implies that the effective population size (N_E_) of this species is ≈ 4.78 × 10^8^ individuals.

*V. fischeri*, in apparent contrast, is well known as a beneficial symbiont of sepiolid squid and monocentrid fishes, in which it produces light that enables the host to evade predators by counter-illumination (Soto et al. 2014). The hosts it colonizes are distributed in different oceans around the world, but it can also be found as a free-living member of the ocean bacterioplankton (Soto et al. 2014). Here also the ecological diversity is borne out by high genetic diversity at a number of loci, with estimated πs = 0.067 and an inferred N_E_ of ≈ 1.62 × 10^8^ (Wollenberg and Ruby 2012). For both species, and for the *Vibrionacae* more generally, what forces lead to such high genetic diversity within species?

Both mutation and horizontal-gene transfer (HGT) are expected to contribute to *Vibrio* biodiversity, but evidence exists that mutation is the primary force driving diversification within *Vibrio* clades (Thompson et al. 2004; Sawabe et al. 2009; Vos and Didelot 2009). It has therefore been tempting to invoke high mutation rates in *Vibrio* species to explain their high genetic diversity. However, we show here that both bpsm and indel rates in *V. fischeri* and *V. cholerae* are low, even for bacteria (Figure S2). In fact, *V. cholerae* has one of the lowest recorded genome-wide rates of bpsms (0.0004 /genome/generation) and indels (0.00007/genome/generation) of any bacterial species (Sung et al. 2016). We suggest that the high genetic diversity and low mutation rates in these *Vibrio* species can be reconciled by the drift-barrier hypothesis, which states generally that any trait, including replication fidelity, may be refined by natural selection only to the point at which further improvement becomes overwhelmed by the power of genetic drift (Lynch 2010; Lynch 2011; Sung et al. 2012; Sung et al. 2016). Natural selection is most powerful in large populations of organisms with genomes composed of a high amount of coding sequence. Although *Vibrio* genomes are not exceptionally large, most sites are coding and their effective population sizes are among the highest recorded (Wollenberg and Ruby 2012; Sung et al. 2016). Thus, both high amounts of coding sequence and high effective population size increase the ability of natural selection to reduce both bpsm and indel rates (Lynch 2010; Lynch 2011; Sung et al. 2012), yet yield enormous allelic diversity at any given time in both of these *Vibrio* species (Thompson et al. 2004; Vos and Didelot 2009).

If the extremely low mutation rates of these *Vibrio* species are a product of powerful selection enabled by massive populations to refine the machinery of DNA replication, it follows that many other heritable traits may have also been optimized in these organisms. One of the best known *Vibrio* traits is their extremely rapid growth rates, with several species achieving doubling times of 10–15 minutes (Brenner et al. 2005). Growth rate is a broad polygenic character requiring the coordination of essentially all systems, including ribosome synthesis, catabolic and anabolic pathways, and of course, replication. *Vibrio* species are also typified by having genomes with two chromosomes, which provide two origins of replication that could conceivably accelerate DNA replication. This genome architecture has clearly been refined by selection for rapid growth: not only does the experimental reduction of the *V. cholerae* genome from two chromosomes to one decrease growth rates by 25–40% (Val et al. 2012), the early-replicating chr1 is enriched relative to chr2 in genes required for rapid growth relative (Cooper et al. 2010). Other *Vibrio* traits (metabolism, regulatory capacity in fluctuating environments, stress tolerance) in addition to their high-fidelity replication and rapid growth may also conceivably have been exceptionally refined by selection. Put more broadly, the drift-barrier hypothesis may be extended and applied to many traits, rendering the prediction that DNA replication fidelity and the optimality of many other phenotypes may be positively correlated – the lower the mutation rate, the more selectively optimized the polygenic trait.

### Mutation rate and spectra variation among genome regions

This hypothesis that *Vibrio* replication has been exquisitely optimized relative to other bacterial genomes because of their high coding content and high effective population sizes does not mean that all regions of *Vibrio* genomes have been equivalently refined relative to one another. Both bpsm and indel rates and spectra varied among genomic regions of *V. fischeri* and *V. cholerae*, which may systematically affect genome composition. In the *Vf*-wt lines, both G:C > A:T transitions and G:C > T:A transversions occurred at higher rates on chr2, although only the rate of G:C > T:A transversions was significantly higher (Figure 1C). Using experiment-wide estimates of each conditional bpsm rate in *V. fischeri*, we estimate that the %GC-content should be 0.184 in the absence of selection and recombination. Yet, the %GC-content of chr2 is expected to be 0.156 at mutation-drift equilibrium, 0.047 lower than expectations for chr1 (0.203). The actual %GC content of chr2 of the *V. fischeri* ES114 genome is also lower than chr1 (chr1: 0.390; chr2: 0.370) suggesting that bpsm biases on these chromosomes have contributed to this pattern. Even stronger biases differentiating the chromosomes are seen in the *Vc*-wt lines, driven by significantly higher A:T > G:C transition rates on chr1 and by non-significant increases in G:C > A:T and G:C > T:A rates on chr2 (Figure 1C). These spectra predict %GC-contents of 0.293 for chr1 and 0.201 for chr2 at mutation-drift equilibrium and likely contribute to the lower realized %GC content of chr2 in *V. cholerae* 2740–80 (chr1: 0.479; chr2: 0.468). Overall, these findings suggest that bpsm pressures contribute to genome-wide and intra-genome variation in %GC contents, but indel biases (Dillon et al. 2015), selection (Hershberg and Petrov 2010; Hildebrand et al. 2010), and/or biased gene conversion (Hershberg and Petrov 2010; Lassalle et al. 2015) must also contribute to the realized %GC contents in *V. fischeri* and *V. cholerae*.

Prior MA studies have reported that the majority of indels involve the loss or gain of a single nucleotide and occur predominantly in simple-sequence repeats (SSRs), where the number of repeated units scales positively with the indel rate (Xu et al. 2000; Lee et al. 2012; Long et al. 2014; Dettman et al. 2016). These SSRs have gained attention not only because they vary sufficiently in repeat number among strains to enable rapid genotyping (van Belkum et al. 1998; Danin-Poleg et al. 2007; Ghosh et al. 2008), but also because in some species they associate with variable heritable expression of genes related to host colonization and disease (Moxon et al. 1994; Field et al. 1999; Moxon et al. 2006), begging the question of whether these mutation-prone sequences have indirectly evolved to enable this plasticity.

In *V. fischeri*, the insertion-deletion rates in SSRs correlated positively with both the number of repeated units and the length of the repeated unit (Figure 2). While the former bias is consistent with a number of previous studies (Xu et al. 2000; Lee et al. 2012; Long et al. 2014; Dettman et al. 2016), the positive correlation between the length of the repeated unit in an SSR and its indel rate has not been reported in previous MA-WGS experiments. One possible reason for this discrepancy might be an increased occurrence of SSRs with longer repeats in the *V. fischeri* ES114 genome (Ruby et al. 2005), which we find to be highly mutagenic (Figure 3). There are 100 SSRs of three or more units in the *V. fischeri* ES114 genome where the repeated unit is at least 4-bps in length. A second possibility is that larger indels have gone undetected by prior MA-WGS analyses focused on MMR-deficient strains, in which single-nucleotide indels are evidently more common (Supplementary Dataset 2). Further, longer indels, especially those in SSRs, are subject to increased false-negative rates due to limitations in the ability of short-read sequencing to resolve them. The majority of multi-nucleotide indels that we identified were supported with very low consensus in the initial alignments because of reads that only partly covered the SSR. Only when we filtered out reads that were not anchored by bps on both sides of the SSR did we achieve high consensus for these indels (Supplementary Dataset 2). It will be interesting to apply these sensitive detection methods for long SSR-associated indels to future experiments to see whether other species also experience elevated indel rates in SSRs with longer repeat units. We emphasize that this experiment also demonstrates that the loss of MMR shifts the spectrum of indel mutations from multi-nucleotide indels in SSRs with longer repeated units towards single nucleotides in homopolymeric runs, a shift with potentially broad phenotypic consequences.

Our study adds to the theory that SSRs may generate localized hyper-mutation in coding regions that may serve as “contingency loci,” enabling potentially beneficial population-level variation in both function and expression of the affected genes (van Belkum et al. 1998; Moxon et al. 2006). These experiments, particularly those conducted in a *ΔmutS* background, identified clusters of frameshift mutations found within genes that could yield adaptive mutant phenotypes in *Vibrio* populations (Table 2). Some in fact have previously been reported to be hypermutable in experimental studies of bacterial function. In *V. cholerae, tcpH* is an important co-regulator that links temperature and pH to control expression of the major virulence regulon (Carroll et al. 1997). During experimental infections of the rabbit model, *tcpH* frameshift mutations occurred at a frequency of 10^−4^ owing to mutations in the same poly-G tract that accumulated in our experiment and they enhanced fitness in culture, which the authors speculated could reflect a benefit during dispersal from the host into the environment (Carroll et al. 1997). Another locus, *mshQ*, encodes a pilus-associated adhesion protein, in which mutants have reported to contribute to the rapid transition between opaque and translucent colonies in *Vibrio* species (Enos-Berlage et al. 2005). These colony types associate with different dynamics of biofilm formation in the laboratory and may reflect different capacity for attachment to natural substrates or other cells, facilitating biofilm differentiation (Yildiz and Visick 2009). Yet another mutagenic homopolymeric tract potentially generating population heterogeneity in biofilm production was found in *vpsT* in *V. fisheri*, a transcriptional activator that regulates polysaccharide production (Yildiz and Visick 2009). Other hypermutable mononucleotide tracts were observed in 14 other loci in the *V. cholerae* and *V. fischeri ΔmutS* lines and were enriched in genes responsible for biofilm production, including putative diguanylate cyclases that affect the regulation of biofilm traits via the messenger molecule cyclic diguanylate and (Yildiz and Visick 2009). Taken together, this study shows that despite inherently very low mutation rates, *Vibrio* populations likely harbor genetic variation at SSR’s that affect traits in which plasticity may be beneficial – such as virulence or adherence mechanisms – and this is especially the case for populations with defective mismatch repair.

**Table 2.**
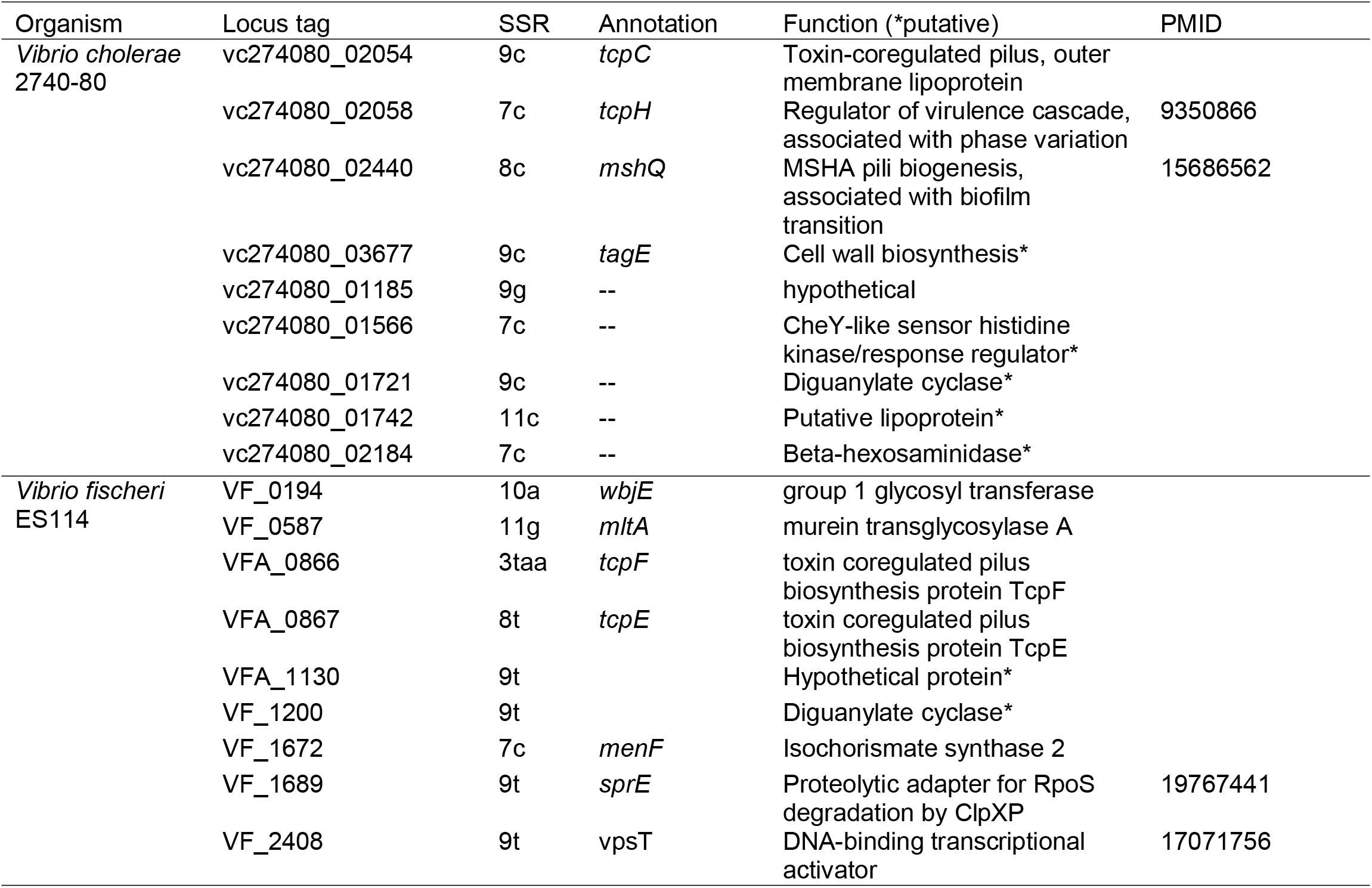
Simple sequence repeats (SSRs) in *Vibrio cholerae* and *Vibrio fischeri* found in coding regions to be mutagenic and associated with gene products whose expression could benefit from population-level plasticity.

### The role of mismatch repair in genome evolution

The mismatch repair pathway is a primary DNA repair pathway in organisms across the tree of life (Kunkel and Erie 2005), but strains lacking MMR are common in nature (Hazen et al. 2009), chronic infections (Hall and Henderson-Begg 2006; Mena et al. 2008; Oliver 2010; Marvig et al. 2013), or long-term evolution experiments (Sniegowski et al. 1997). Loss of a functional MMR system can elevate mutation rates anywhere from 5 to 1000-fold, depending on both the defective component of the pathway and the genetic background (Lyer et al. 2006; Long et al. 2015; Reyes et al. 2015). The primary proteins involved in the MMR pathway in bacteria include the MutS protein, which binds mismatches to initiate repair, the MutL protein, which coordinates multiple steps of MMR synthesis, and the MutH protein, which nicks the unmethylated strand to remove the replication error (Kunkel and Erie 2005). The removal of the *mutS* gene in this study resulted in a 317-fold increase in the bpsm rate and a 102-fold increase in the indel rate in *V. fischeri* ES114. The removal of the *mutS* gene in *V. cholerae* had a less dramatic effect on the bpsm rate (85-fold increase), but a more dramatic effect on the indel rate (142-fold increase). Overall, this suggests that MMR is more central to the repair of bpsms in *V. fischeri*, while it is more important for the repair of indels in *V. cholerae*.

Despite the relatively wide range in the consequences of losing a functional MMR system for bpsm rates, the resulting bpsm spectra were remarkably similar in *V. fischeri* and *V. cholerae*. Specifically, unlike the wild-type experiments, the bpsm spectra of the *Vf*-mut and *Vc*-mut MA experiments were not significantly different, as both were dominated by G:C > A:T and A:T > G:C transitions. Furthermore, the vast majority of indels in the both the *Vf*-mut and *Vc*-mut experiments involve only a single-nucleotide, most of which occurred in homopolymers, where their rates scaled positively with the length of the homopolymer (Figure 5). These observations are consistent with previous reports in other bacterial MA experiments using MMR-deficient strains (Lee et al. 2012; Long et al. 2014; Sung et al. 2015; Dettman et al. 2016), but reveal distinct mutation biases from genotypes with functional MMR. In fact, the strong site-specific biases in the mutation spectra generated by the loss of MMR (Table 2) may help to explain the prevalence of mutator alleles in environmental and clinical settings, despite the inevitable fitness costs of the mutational load. Couce et al. have found that mutator alleles can modify the distribution of fitness effects of individual beneficial mutations by enriching a specific spectrum of spontaneous mutations, and impact the evolutionary trajectories of different strains (Couce et al. 2013; Couce et al. 2015), although the generality of how mutators influence adaptive dynamics requires further study.

The loss of MMR also helps reveal mutation biases associated with the replicative polymerase that cannot be observed using the low number of mutations generated in wild-type MA-WGS experiments (Lee et al. 2012; Sung et al. 2015; Dettman et al. 2016). A mirrored wave-like pattern of bpsm rates on opposing replichores has now been observed in multiple MMR-deficient species studied by MA-WGS, although the exact shape and strength of the pattern varies between species (Foster et al. 2013; Long et al. 2014; Dettman et al. 2016). We find the same mirrored wave-like pattern of bpsm rates on the opposing replichores of chr1 in MMR-deficient *V. fischeri* and *V. cholerae* (Figure 6A, B) and show that bpsm rates of concurrently replicating regions of this chromosome are significantly correlated in both genomes. This suggests that bpsm rates are impacted by genome location and that regions replicated at similar times on opposing replichores experience similar bpsm rates, at least in MMR-deficient strains. However, we do not observe any significant variation in the bpsm rates on chr2 (Figure 6A, B). We suggest that bpsm rates are less variable on chr2 because of their delayed replication. Specifically, chr2 replication is not initiated until a large portion of chr1 has already been replicated (Egan and Waldor 2003; Duigou et al. 2006; Rasmussen et al. 2007), which means that chr2 is not replicated during the primary peaks in the bpsm rate on the opposing replichores of chr1, and thus experience more consistent bpsm rates across the chromosome. These patterns of variation in mutation rates among genome regions suggest that it is conceivable that selection may act to position some genes to avoid regions of higher mutation pressure, especially in populations of the size and diversity of *Vibrio* species.

### Concluding Remarks

Mutation-accumulation experiments paired with whole-genome sequencing enable an unprecedented view of genome-wide mutation rates and spectra and reveal the underlying biases of spontaneous mutation. These underlying biases can explain why some genome regions evolve more rapidly than others, why the coding content of different genome regions varies, and perhaps how clonal populations may generate adaptively diverse progeny. The primary properties of *V. fischeri* and *V. cholerae* spontaneous mutation that we have identified in this MA-WGS study are: (a) base-substitution and insertion-deletion mutation rates are low, consistent with other bacterial species; (b) base-substitution mutation biases vary between chromosomes, but don’t fully explain their realized %GC contents; (c) both the length of repeat units and the number of repeated units in simple-sequence repeats correlate positively with the insertion-deletion rate of the SSR; (d) loss of a proficient mismatch repair system has inconsistent effects on base-substitution and indel mutation rates in different taxa, but consistently generates convergent mutation spectra that are dominated by transitions and short indels; and (e) base-substitution mutations in strains deficient in mismatch repair vary in a mirrored wave-like pattern on opposing replichores on chromosome 1, but variation is limited on chromosome 2. As we uncover properties of spontaneous mutation in diverse microbes, we can continue to assess the generality of mutational biases and more accurately evaluate the role of mutation bias in molecular evolution.

## MATERIALS AND METHODS

### Bacterial strains and culture conditions

The two wild-type MA experiments were founded from a single clone derived from *V. fischeri* ES114 and *V. cholerae* 2740–80, respectively. All MA experiments with *V. fischeri* were carried out on tryptic soy agar plates supplemented with NaCl (TSAN) (30 g/liter tryptic soy broth powder, 20 g/liter NaCl, 15 g/liter agar) and were incubated at 28°. Frozen stocks of each MA lineage were prepared at the end of the experiment by growing a single colony overnight in 5ml of tryptic soy broth supplemented with NaCl (TSBN) (30 g/liter tryptic soy broth powder, 20 g/liter NaCl) at 28° and freezing in 8% DMSO at −80°. For *V. cholerae*, all MA experiments were carried out on tryptic soy agar plates (TSA) (30 g/liter tryptic soy broth powder, 15 g/liter agar) and were incubated at 37°. Similarly, frozen stocks were prepared by growing a single colony from each lineage overnight in 5ml of tryptic soy broth (TSB) (30 g/liter tryptic soy broth powder) at 37° and were stored in 8% DMSO at - 80°.

Mutator strains of *V. fischeri* ES114 and *V. cholerae* 2740–80 were generated by replacing the *mutS* gene in each genome with an erythromycin resistance cassette, as described previously (Datsenko and Wanner 2000; Heckman and Pease 2007; Val et al. 2012). Briefly, we used splicing by overlap extension (PCR-SOE) to generate two erythromycin resistance cassettes, one of which was flanked by ≈ 750 bps of the upstream and downstream regions of the *mutS* gene in *V. fischeri* ES114, while the second was flanked by ≈ 750 bps of the upstream and downstream regions of the *mutS* gene in *V. cholerae* 2740–80 (Heckman and Pease 2007). Both the *V. fischeri* ES114 and *V. cholerae* 2740–80 *ΔmutS* fragments were then cloned into the R6K γ-ori-based suicide vector pSW7848, which contains a *ccdB* toxin gene that is arabinose-inducible and glucose-repressible *(P_BAD_*) (Val et al. 2012). Both of these pSW7848 plasmids, henceforth referred to as pSW7848-*VfΔmutS* and pSW7848-*VcΔmutS*, were transformed into *Escherichia coli* pi3813 chemically competent cells and stored at −80° (Datsenko and Wanner 2000).

Conjugal transfer of the pSW7848-*VfΔmutS* and pSW7848-*VcΔmutS* plasmids was performed using a tri-parental mating with the *E. coli* pi3813 cells as the donors (Val et al. 2012), *E. coli* DH5α-pEVS104 as the helper (Stabb and Ruby 2002), and *V. fischeri* ES114 and *V. cholerae* 2740–80 as the respective recipients. For *V. fischeri* ES114, the chromosomally inserted pSW7848-*VcΔmutS* plasmid resulting from a crossover at the *ΔmutS* gene was selected on LBS plates (Graf et al. 1994) containing 1% glucose and 1ug/ml chloramphenicol at 28°. Selection for loss of the plasmid backbone from a second recombination step was then performed on LBS plates containing 0.2% arabinose at 28°, which induces the *P_BAD_* promoter of the *ccdB* gene and ensures that all cells that have not lost the integrated plasmid will die (Val et al. 2012). For *V. cholerae*, the chromosomally inserted pSW7848-*VcΔmutS* plasmid was selected on LB plates (Sambrook et al. 1989) containing 1% glucose and 5ug/ml chloramphenicol at 30°. Selection for loss of the plasmid backbone was performed on LB plates with 0.2% arabinose at 30°. Replacement of the *mutS* gene in *V. fischeri* ES114 and *V. cholerae* 2740–80 were verified by conventional sequencing, and *V. fischeri* ES114 *ΔmutS* and *V. cholerae* 2740–80 *ΔmutS* were used to found the two mutator MA experiments, under identical conditions to those described above for the wild-type experiments.

### Ancestral reference genomes

Prior to this study, the genome of *V. fischeri* ES114 was already in completed form and annotated, consisting of three contigs representing chr1, chr2, and the 45.85 Kb plasmid (Ruby et al. 2005). Further, the location of the *oriC* on both chromosomes was available in dOriC 5.0, a database for the predicted *oriC* regions in bacterial and archaeal genomes (Gao et al. 2013). Fortunately, the *oriC* region on both chromosomes had been placed at coordinate zero, allowing us to proceed with this *V. fischeri* ES114 reference genome for all subsequent *V. fischeri* analyses. In contrast, when we initiated our MA experiment the *V. cholerae* 2740–80 genome was still in draft form, consisting of 257 scaffolds with unknown chromosome association. Therefore, to reveal inter-chromosomal variation and assess the effects of genome location on bpsm and indel rates, we used single molecule, real-time (SMRT) sequencing to generate a complete assembly separated into the two contigs of *V. cholerae* 2740–80.

The Pacific Biosciences RSII sequencer facilitates the completion of microbial genomes by producing reads of multiple kilobases (kb) that extend across repetitive regions and allow whole-genomes to be assembled at a relatively limited cost (Koren and Phillippy 2015). Genomic DNA (gDNA) was prepared using the Qiagen Genomic-Tip Kit (20/G) from overnight cultures of *V. cholerae* 2740–80 grown in LB at 37° using manufacturer’s instructions. Importantly, this kit uses gravity filtration to purify gDNA, which limits shearing and increases the average fragment size of the resulting gDNA sample. Long insert library preparation and SMRT sequencing was performed on this *V. cholerae* 2740<80 gDNA at the Icahn School of Medicine at Mount Sinai according to the manufacturer’s instructions, as described previously (Beaulaurier et al. 2015). Briefly, libraries were size selected using Sage Science Blue Pippin 0.75% agarose cassettes to enrich for long-reads, and were assessed for quantity and insert size using an Agilent DNA 12,000 gel chip. Primers, polymerases, and magnetic beads were loaded to generate a completed SMRTbell library, which was run in a single SMRT cell of a Pacific Biosciences RSII sequencer at a concentration of 75 pM for 180 minutes.

As expected, our long insert SMRT sequencing library generated mostly long reads, with an average sub-read length of 8,401 bps and an N50 of 11,480 bps. We used the hierarchical genome-assembly process workflow (HGAP3) to generate a completed assembly of *V. cholerae* 2740–80 and polished our assembly using the Quiver algorithm (Chin et al. 2013). The resultant assembly consisted of two contigs representing chr1 and chr2, with an average coverage of 128x. We annotated this assembly using prokka (v1.11), specifying *Vibrio* as the genus (Seemann 2014). We then identified the location of the *oriC* on both contigs using Ori-finder, which applies analogous methods to those used by dOriC 5.0 to identify *oriC* regions in bacterial genomes (Gao and Zhang 2008; Gao et al. 2013). Of course, these *oriC* regions were not located at coordinate zero of the *V. cholerae* 2740–80 reference genome, so we reformatted the reference genome to place each *oriC* region at the beginning of the chr1 and chr2 contigs, then stitched the contigs back together and re-polished the genome using Quiver. Prokka was then run a second time to update the location of all genes, and this re-formatted *V. cholerae* 2740–80 genome was used as the ancestral reference genome for all subsequent *V. cholerae* analyses.

### MA-WGS Process

For the two wild-type MA experiments, seventy-five independent lineages were founded by single cells derived from a single colony of *V. fischeri* ES114 and *V. cholerae* 2740–80, respectively. Each of these lineages was then independently propagated every 24 hours onto fresh TSAN for *V. fischeri* and fresh TSA for *V. cholerae*. This cycle was then repeated for a total of 217 days. For the two mutator MA experiments, forty-eight independent lineages were founded and propagated as described above from a single colony each of *V. fischeri* ES114 *ΔmutS* and *V. cholerae* 2740–80 *ΔmutS*, respectively. However, because of their higher mutation rates, these lineages were only propagated for a total of 43 days. At the conclusion of the four MA experiments, each lineage was grown overnight in the appropriate liquid broth at the appropriate temperature (see above), and stored at −80° in 8% DMSO.

Daily generations were estimated monthly for the wild-type lineages and bi-monthly for the mutator lineages by calculating the number of viable cells in a representative colony from 10 lineages per MA experiment following 24 hours of growth. During each measurement, the representative colonies were placed in 2 ml of phosphate buffer saline (80 g/liter NaCl, 2 g/liter KCl, 14.4 g/liter Na_2_HPO_4_ • 2H_2_O, 2.4 g//liter KH2PO4), serially diluted, and spread plated on TSAN or TSA for *V. fischeri* and *V. cholerae*, respectively. These plates were then incubated for 24 hours at 28° or 37°, and the daily generations per colony were calculated from the number of viable cells in each representative colony. The average daily generations were then calculated for each time-point using the ten representative colonies, and the total generations elapsed between each measurement were calculated as the product of the average daily generations and the number of days before the next measurement. The total of number of generations elapsed during the duration of the MA experiment per lineage was then calculated as the sum of these totals over the course of each MA study (Figure S1).

At the conclusion of each of the four MA experiments, gDNA was extracted using the Wizard Genomic DNA Purification Kit (Promega) from 1 ml of overnight culture (TSBN at 28° for *V. fischeri*; TSB at 37° for *V. cholerae)* inoculated from 50 representative stored lineages for *Vf*-wt and *Vc*-wt experiments, and all 48 stored lineages for the *Vf*-mut and *Vc*-mut experiments. For the wild-type MA experiments, gDNA from the ancestral *V. fischeri* ES114 and *V. cholerae* 2740–80 strains was also extracted. All libraries were prepared using a modified Illumina Nextera protocol designed for inexpensive library preparation of microbial genomes (Baym et al. 2015). Sequencing of the *Vf*-wt and *Vc*-wt lineages and their respective ancestors was performed using the 101-bp paired-end Illumina HiSeq platform at the Beijing Genome Institute (BGI), while sequencing of the *Vf*-mut and *Vc*-mut lineages was performed using the 151-bp paired-end Illumina HISeq platform at the University of New Hampshire Hubbard Center for Genomic Studies.

Our raw fastQ reads were analyzed using fastQC, and revealed that 48 *Vf*-wt lineages, 49 *Vc*-wt lineages, 19 *Vf*-mut lineages, and 22 *Vc*-mut lineages were sequenced at sufficient depth to accurately identify bpsm and indel mutations. Our failure to successfully sequence a high proportion of *Vf*-mut and *Vc*-mut lineages was mostly generated by a poorly normalized library, leading to limited sequence data for several of the mutator lineages. For the successfully sequenced lineages, all reads were mapped to their respective reference genomes with both the Burrows-Wheeler Aligner (BWA) (Li and Durbin 2009) and Novoalign (www.novocraft.com). The average depth of coverage across the successfully sequenced lineages of each MA experiment was 100x for *Vf*-wt, 96x for *Vc*-wt, 124x for *Vf*-mut, and 92x for *Vc*-mut.

### Base-substitution mutation identification

For all four MA experiments, bpsms were identified as described previously in an MA experiment with *Burkholderia cenocepacia* (Dillon et al. 2015). Briefly, we used SAMtools to convert the SAM alignment files from each lineage to mpileup format (Li et al. 2009), then in-house perl scripts to produce the forward and reverse read alignments for each position in each line. A three-step process was then used to detect putative bpsms. First, pooled reads across all lines were used to generate an ancestral consensus base at each site in the reference genome. This allows us to correct for any differences that may exist between the reference genomes and the ancestral colony of each our MA experiments. Second, a lineage specific consensus base was generated at each site in the reference genome for each individual MA lineage using only the reads from that line. Here, a lineage specific consensus base was only called if the site was covered by at least two forward and two reverse reads and at least 80% of the reads identified the same base. Otherwise, the site was not analyzed. Third, each lineage specific consensus base that was called was compared to the overall ancestral consensus of the MA experiment and a putative bpsm was identified if they differed. This analysis was carried out independently with the alignments generated by BWA and Novoalign, and putative bpsms were considered genuine only if both pipelines independently identified the bpsm and they were only identified in a single lineage.

Using relatively lenient criteria for identifying lineage specific consensus bases, we were able to analyze the majority of the genome in all of our lineages, but increase our risk of falsely identifying bpsms at low coverage sites. Therefore, we generated a supplementary dataset for all genuine bpsms identified in this study, which includes the read coverage and consensus at each site where a bpsm was identified (Supplementary Dataset 1). We do not see clusters of bpsms at the lower limits of our coverage or consensus requirements. In fact, the vast majority of bpsms in all of four MA experiments were covered by more than 50 reads and were supported by more than 95% of the reads that covered the site. Furthermore, we verified that none of the bpsms that we identified were present in the ancestral *V. fischeri* ES114 and *V. cholerae* 2740–80 strains that we sequenced, so we are confident that nearly all of the bpsms identified in this study were spontaneous bpsms that arose during the mutation accumulation experiments.

### Insertion-deletion mutation identification

All indels identified in this study were also identified using similar requirements to those described in our MA experiment with *B. cenocepacia* (Dillon et al. 2015), with a few modifications. First, we extracted all putative indels from the BWA and Novoalign alignments for each lineage under the requirements that the indel was covered by at least two forward and two reverse reads, and 30% of those reads identified the exact same indel (size and motif). All putative indels that were independently identified by both BWA and Novoalign, where 80% of the reads identified the exact same indel were considered genuine. Next, we noticed that nearly all putative indels that had been identified with less than 80% consensus were in SSRs. Therefore, for indels where only 30–80% of the reads identified the exact same indel, we parsed out only reads that had bases on both the upstream and downstream region of the SSR (if the indel was in an SSR), and on both the upstream and downstream region of the indel itself (if the indel was not in an SSR). Using only this subset of reads, we reassessed the number of reads that identified the exact same indel (size and motif), and considered these initially low confidence putative indels genuine if more than 80% of these sub-reads identified the exact same indel. In addition, we passaged our alignment output through the pattern-growth algorithm PINDEL to identify any large-scale genuine indels using paired-end information (Ye et al. 2009). Here, we required a total of 20 reads, with at least 6 forward and 6 reverse reads, and 80% of the reads to identify the exact same indel for the indel to be considered genuine. This summative collection of genuine indels was then compared to the analysis of the ancestral *V. fischeri* ES114 and *V. cholerae* 2740–80 strains, and any indels that were identified in their corresponding ancestor, or more than 50% of the analyzed lineages from the corresponding MA experiment were excluded from subsequent analyses.

Our initial filters for indels were even more lenient than those for bpsms, which could have led to false positive indel identification in the putative indel phase. However, we subsequently required at least 80% consensus for all genuine indels identified in this study among reads that had bases covering both the upstream and downstream regions of putative indels that were not in an SSR. For genuine indels that were in an SSR, we required at least 80% consensus among reads that had bases covering both the upstream and downstream regions of the SSR. Further, we verified that all indels identified were not present in our ancestral *V. fischeri* ES114 or *V. cholerae* 2740–80 strains and were not identified in more than 50% of the other lineages analyzed in the same MA experiment. As with our bpsms, we generated a supplementary dataset containing all genuine indels analyzed in this study, which includes read coverage and consensus of each site where an indel was identified, as well as the read coverage and consensus among reads with bases covering both the upstream and downstream regions of the indel or SSR if the initial consensus among reads covering the indel was below 80% (Supplementary Dataset 2). Thus, we are confident that nearly all indels identified in this study were genuine spontaneous indels that arose during the mutation accumulation process.

### Mutation-rate analyses

Overall bpsm and indel rates were calculated for each lineage using the equation: *μ* = *m/nT*, where *μ* represents the mutation rate, represents the number of mutations observed, *n* represents the number of ancestral sites analyzed, and *T* represents the total number of generations elapsed per lineage. Conditional bpsm rates for each lineage were calculated using the same equation, but with *m* representing the number of bpsms of the focal bpsm type, and *n* representing the number analyzed ancestral sites that could generate the focal bpsm type. All summative bpsm and indel rates presented for each MA experiment were calculated as the average mutation rate across all analyzed lineages, while summative standard errors were calculated as the standard deviation of the mutation rate across all lines (*s*), divided by the square root of the total number of lines in the corresponding MA experiment 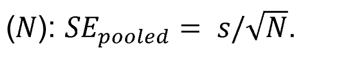.

For our analysis of bpsm and indel rates within chromosomes, we divided each chromosome into 100 kb intervals, starting at the origin of replication and extending bi-directionally to the replication terminus. Bpsm rates in each interval were measured by dividing the total number of bpsm or indel mutations from this study, by the product of the total number of sites analyzed in each interval across all lines and the number of generations per line, using the same formula described above for genome wide mutation rates: *μ* = *m/nT*. Because none of the chromosomes were exactly divisible by 100 kb, the final intervals on each replichore were always less than 100 kb, but their mutation rates were calibrated to the number of of bases analyzed.

Potential effects of selection on observed mutations were assessed as follows. For each MA experiment, the expected ratio of coding to non-coding mutations was determined directly from each ancestral reference genome, and the expected ratio of synonymous to nonsynonymous bpsms was calculated from each ancestral reference genome, after accounting for codon usage and %GC content of synonymous and nonsynonymous sites.

### Data availability

Separate Bioprojects have been generated on NCBI for all sequencing data that pertains to these *V. fischeri* (PRJNA256340) and *V. cholerae* (PRJNA256339) MA-WGS projects. Under these Bioprojects, Illumina DNA sequences for all the *V. fischeri* and *V. cholerae* MA-WGS lines are available under the following Biosamples: *Vf*-wt = SAMN05366916, *Vc*-wt = SAMN05366910, *Vf*-mut = SAMN05366917, *Vc*-mut = SAMN05366911. The completed V. *cholerae* 2740–80 genome assembly generated in this study is also available under the Biosample SAMN05323685. All strains are available upon request.

### Statistical analyses

All statistical analyses were performed in R Studio Version 0.99.489 using the Stats analysis package (R Development Core Team 2013).

## ACKNOWLEDGMENTS

We thank Cheryl Whistler and Randi Foxall for their input on creating knockout mutants in *Vibrio* species, and Brian VanDam for technical support. This work was supported by the Multidisciplinary University Research Initiative Award from the US Army Research Office (W911NF-09-1-0444 to ML, P. Foster, H. Tang, and S. Finkel); and the National Science Foundation Career Award (DEB-0845851 to VSC).

## REFRENCES

Agier N, Fischer G. 2012. The mutational profile of the yeast genome is shaped by replication. Mol. Biol. Evol. 29:905–913.

Baym M, Kryazhimskiy S, Lieberman TD, Chung H, Desai MM, Kishony R. 2015. Inexpensive multiplexed library preparation for megabase-sized genomes. PLoS One 10:e0128036.

Beaulaurier J, Zhang X, Zhu S, Sebra R, Rosenbluh C, Deikus G, Shen N, Munera D, Waldor MK, Chess A, et al. 2015. Single molecule-level detection and long read-based phasing of epigenetic variations in bacterial methylomes. Nat. Commun. 6:7438.

Behringer MG, Hall DW. 2016. The repeatability of genome-wide mutation rate and spectrum estimates. Curr. Genet. 1:1–6.

van Belkum A, Scherer S, van Alphen L, Verbrugh H. 1998. Short-sequence DNA repeats in prokaryotic genomes. Microbiol. Mol. Biol. Rev. 62:275–293.

Brenner DJ, Krieg NR, Staley JT, Garrity GM. 2005. Bergey’s Manual of Systematic Bacteriology. 2nd ed. New York: Springer-Verlag

Carroll P a, Tashima KT, Rogers MB, DiRita VJ, Calderwood SB. 1997. Phase variation in *tcpH* modulates expression of the ToxR regulon in *Vibrio cholerae*. Mol. Microbiol. 25:1099–1111.

Chen C-L, Rappailles A, Duquenne L, Huvet M, Guilbaud G, Farinelli L, Audit B, D’Aubenton-Carafa Y, Arneodo A, Hyrien O, et al. 2010. Impact of replication timing on non-CpG and CpG substitution rates in mammalian genomes. Genome Res. 20:447–457.

Chin C-S, Alexander DH, Marks P, Klammer AA, Drake J, Heiner C, Clum A, Copeland A, Huddleston J, Eichler EE, et al. 2013. Nonhybrid, finished microbial genome assemblies from long-read SMRT sequencing data. Nat. Methods 10:563–569.

Cooper VS, Vohr SH, Wrocklage SC, Hatcher PJ. 2010. Why genes evolve faster on secondary chromosomes in bacteria. Plos Comput. Biol. 6:e1000732.

Couce A, Guelfo J, Blazquez J. 2013. Mutational spectrum drives the rise of mutator bacteria. Plos Genet. 9:e1003167.

Couce A, Rodriguez-Rojas A, Blazquez J. 2015. Bypass of genetic constraints during mutator evolution to antibiotic resistance. Proc. R. Soc. London Ser. B-Biological Sci. 282:20142698.

Danin-Poleg Y, Cohen LA, Gancz H, Broza YY, Goldshmidt H, Malul E, Valinsky L, Lerner L, Broza M, Kashi Y. 2007. *Vibrio cholerae* strain typing and phylogeny study based on simple sequence repeats. J. Clin. Microbiol. 45:736–746.

Datsenko KA, Wanner BL. 2000. One-step inactivation of chromosomal genes in *Escherichia coli* K-12 using PCR products. Proc. Natl. Acad. Sci. U. S. A. 97:6640–6645.

Denver DR, Dolan PC, Wilhelm LJ, Sung W, Lucas-Lledo JI, Howe DK, Lewis SC, Okamoto K, Thomas WK, Lynch M, et al. 2009. A genome-wide view of *Caenorhabditis elegans* base-substitution mutation processes. Proc. Natl. Acad. Sci. U. S. A. 106:16310–16314.

Dettman JR, Sztepanacz JL, Kassen R. 2016. The properties of spontaneous mutations in the opportunistic pathogen *Pseudomonas aeruginosa*. BMC Genomics 17:27–41.

Dillon MM, Sung W, Lynch M, Cooper VS. 2015. The rate and molecular spectrum of spontaneous mutations in the GC-rich multichromosome genome of *Burkholderia cenocepacia*. Genetics 200:935–946.

Duigou S, Knudsen KG, Skovgaard O, Egan ES, Lobner-Olesen A, Waldor MK. 2006. Independent control of replication initiation of the two *Vibrio cholerae* chromosomes by DnaA and RctB. J. Bacteriol. 188:6419–6424.

Egan ES, Waldor MK. 2003. Distinct replication requirements for the two *Vibrio cholerae* chromosomes. Cell 114:521–530.

Enos-Berlage JL, Guvener ZT, Keenan CE, McCarter LL. 2005. Genetic determinants of biofilm development of opaque and translucent *Vibrio parahaemolyticus*. Mol. Microbiol. 55:1160–1182.

Field D, Magnasco MO, Moxon ER, Metzgar D, Tanaka MM, Wills C, Thaler DS. 1999. Contingency loci, mutator alleles, and their interactions: Synergistic strategies for microbial evolution and adpatation in pathogenesis. Mol. Strateg. Biol. Evol. 870:378–382.

Foster PL, Hanson AJ, Lee H, Popodi EM, Tang HX. 2013. On the mutational topology of the bacterial genome. G3-Genes Genomes Genet. 3:399–407.

Gao F, Luo H, Zhang CT. 2013. DoriC 5.0: an updated database of *oriC* regions in both bacterial and archaeal genomes. Nucleic Acids Res. 41:D90–D93.

Gao F, Zhang C-T. 2008. Ori-Finder: a web-based system for finding *oriCs* in unannotated bacterial genomes. BMC Bioinformatics 9:79–85.

Ghosh R, Nair GB, Tang L, Morris JG, Sharma NC, Ballal M, Garg P, Ramamurthy T, Stine OC. 2008. Epidemiological study of *Vibrio cholerae* using variable number of tandem repeats. FEMS Microbiol. Lett. 288:196–201.

Goldberg S, Murphy JR. 1983. Molecular epidemiological-studies of United-States gulf-coast *Vibrio cholerae* strains-integration site of mutator *Vibriophage* Vca-3. Infect. Immun. 42:224–230.

Graf J, Dunlap P V., Ruby EG. 1994. Effect of transposon-induced motility mutations on colonization of the host light organ by *Vibrio fischeri*. J. Bacteriol. 176:6986–6991.

Hall LMC, Henderson-Begg SK. 2006. Hypermutable bacteria isolated from humans-a critical analysis. Microbiology 152:2505–2514.

Hazen TH, Kennedy KD, Chen S, Yi S V., Sobecky PA. 2009. Inactivation of mismatch repair increases the diversity of *Vibrio parahaemolyticus*. Environ. Microbiol. 11:1254–1266.

Heckman KL, Pease LR. 2007. Gene splicing and mutagenesis by PCR-driven overlap extension. Nat Protoc 2:924–932.

Heidelberg JF, Eisen JA, Nelson WC, Clayton RA, Gwinn ML, Dodson RJ, Haft DH, Hickey EK, Peterson JD, Umayam L, et al. 2000. DNA sequence of both chromosomes of the cholera pathogen *Vibrio cholerae*. Nature 406:477–483.

Heilbron K, Toll-Riera M, Kojadinovic M, Maclean RC. 2014. Fitness is strongly influenced by rare mutations of large effect in a microbial mutation accumulation experiment. Genetics 197:981–990.

Hershberg R, Petrov DA. 2010. Evidence that mutation is universally biased towards AT in bacteria. PLoS Genet. 6:e1001115.

Hildebrand F, Meyer A, Eyre-Walker A. 2010. Evidence of selection upon genomic GC-content in bacteria. PLoS Genet. 6:e1001107.

Koren S, Phillippy AM. 2015. One chromosome, one contig: complete microbial genomes from long-read sequencing and assembly. Curr. Opin. Microbiol. 23:110–120.

Kunkel TA, Erie DA. 2005. DNA mismatch repair. Annu. Rev. Biochem. 74:681–710.

Lassalle F, Périan S, Bataillon T, Nesme X, Duret L, Daubin V. 2015. GC-content evolution in bacterial genomes: the biased gene conversion hypothesis expands. PLOS Genet. 11:e1004941.

Lee H, Popodi E, Tang HX, Foster PL. 2012. Rate and molecular spectrum of spontaneous mutations in the bacterium *Escherichia coli* as determined by whole-genome sequencing. Proc. Natl. Acad. Sci. U. S. A. 109: E2774–E2783.

Li H, Durbin R. 2009. Fast and accurate short read alignment with Burrows-Wheeler transform. Bioinformatics 25:1754–1760.

Li H, Handsaker B, Wysoker A, Fennell T, Ruan J, Homer N, Marth G, Abecasis G, Durbin R. 2009. The sequence alignment/map format and SAMtools. Bioinformatics 25:2078–2079.

Lind PA, Andersson DI. 2008. Whole-genome mutational biases in bacteria. Proc. Natl. Acad. Sci. U.S. A. 105:17878–17883.

Long H, Kucukyildirim S, Sung W, Williams E, Lee H, Ackerman M, Doak TG, Tang H, Lynch M. 2015. Background mutational features of the radiation-resistant bacterium *Deinococcus radiodurans*. Mol. Biol. Evol. 32:2383–2392.

Long H, Sung W, Miller SF, Ackerman MS, Doak TG, Lynch M. 2014. Mutation rate, spectrum, topology, and context-dependency in the DNA mismatch repair (MMR) deficient *Pseudomonas fluorescens* ATCC948. Genome Biol. Evol. 7:262–271.

Lyer RR, Pluciennik A, Burdett V, Modrich PL. 2006. DNA mismatch repair: Functions and mechanisms. Chem. Rev. 106:302–323.

Lynch M, Sung W, Morris K, Coffey N, Landry CR, Dopman EB, Dickinson WJ, Okamoto K, Kulkarni S, Hartl DL, et al. 2008. A genome-wide view of the spectrum of spontaneous mutations in yeast. Proc. Natl. Acad. Sci. U. S. A. 105:9272–9277.

Lynch M. 2010. Evolution of the mutation rate. Trends Genet. 26:345–352.

Lynch M. 2011. The lower bound to the evolution of mutation rates. Genome Biol. Evol. 3:1107–1118.

Martincorena I, Seshasayee ASN, Luscombe NM. 2012. Evidence of non-random mutation rates suggests an evolutionary risk management strategy. Nature 485:95–98.

Marvig RL, Johansen HK, Molin S, Jelsbak L. 2013. Genome analysis of a transmissible lineage of *Pseudomonas aeruginosa* reveals pathoadaptive mutations and distinct evolutionary paths of hypermutators. PLoS Genet. 9:e1003741.

Mena A, Smith EE, Burns JL, Speert DP, Moskowitz SM, Perez JL, Oliver A. 2008. Genetic adaptation of *Pseudomonas aeruginosa* to the airways of cystic fibrosis patients is catalyzed by hypermutation. J. Bacteriol. 190:7910–7917.

Mira A, Ochman H. 2002. Gene location and bacterial sequence divergence. Mol. Biol. Evol. 19:1350–1358.

Morrow JD, Cooper VS. 2012. Evolutionary effects of translocations in bacterial genomes. Genome Biol. Evol. 4:1256–1262.

Moxon ER, Rainey PB, Nowak MA, Lenski RE. 1994. Adaptive evolution of highly mutable loci in pathogenic bacteria. Curr. Biol. 4:24–33.

Moxon R, Bayliss C, Hood D. 2006. Bacterial contingency loci: the role of simple sequence DNA repeats in bacterial adaptation. Annu. Rev. Genet. 40:307–333.

Oliver A. 2010. Mutators in cystic fibrosis chronic lung infection: Prevalence, mechanisms, and consequences for antimicrobial therapy. Int J Med Microbiol 300:563–572.

Ossowski S, Schneeberger K, Lucas-Lledo JI, Warthmann N, Clark RM, Shaw RG, Weigel D, Lynch M. 2010. The rate and molecular spectrum of spontaneous mutations in *Arabidopsis thaliana*. Science 327:92–94.

R Development Core Team. 2013. R: A Language and Environment for Statistical Computing. Rasmussen T, Jensen RB, Skovgaard O. 2007. The two chromosomes of *Vibrio cholerae* are initiated at different time points in the cell cycle. Embo J. 26:3124–3131.

Reyes GX, Schmidt TT, Kolodner RD, Hombauer H. 2015. New insights into the mechanism of DNA mismatch repair. Chromosoma 124:443–462.

Ruby EG, Urbanowski M, Campbell J, Dunn A, Faini M, Gunsalus R, Lostroh P, Lupp C, McCann J, Millikan D, et al. 2005. Complete genome sequence of *Vibrio fischeri:* A symbiotic bacterium with pathogenic congeners. Proc. Natl. Acad. Sci. U. S. A. 102:3004–3009.

Sambrook J, Fritsch EF, Maniatis T. 1989. Molecular Cloning: A Laboratory Manual. New York

Sawabe T, Koizumi S, Fukui Y, Nakagawa S, Ivanova EP, Kita-Tsukamoto K, Kogure K, Thompson FL. 2009. Mutation is the main driving force in the diversification of the *Vibrio splendidus* clade. Microbes Environ. 24:281–285.

Schoolnik GK, Yildiz FH. 2000. The complete genome sequence of *Vibrio cholerae:* a tale of two chromosomes and of two lifestyles. Genome Biol. 1:reviews1016.1–1016.3.

Schrider DR, Houle D, Lynch M, Hahn MW. 2013. Rates and genomic consequences of spontaneous mutational events in *Drosophila melanogaster*. Genetics 194:937–954.

Seemann T. 2014. Prokka: Rapid prokaryotic genome annotation. Bioinformatics 30:2068–2069.

Sharp PM, Shields DC, Wolfe KH, Li WH. 1989. Chromosomal location and evolutionary rate variation in enterobacterial genes. Science 246:808–810.

Sniegowski PD, Gerrish PJ, Lenski RE. 1997. Evolution of high mutation rates in experimental populations of *E. coli*. Nature 387:703–705.

Soto W, Rivera FM, Nishiguchi MK. 2014. Ecological diversification of *Vibrio fischeri* serially passaged for 500 generations in novel squid host *Euprymna tasmanica*. Microb. Ecol. 67:700–721.

Stabb E V., Ruby EG. 2002. RP4-based plasmids for conjugation between *Escherichia* coli and members of the *Vibrionaceae*. Methods Enzymol. 358:413–426.

Stamatoyannopoulos JA, Adzhubei I, Thurman RE, Kryukov G V, Mirkin SM, Sunyaev SR. 2009. Human mutation rate associated with DNA replication timing. Nat. Genet. 41:393–395.

Stokke C, Waldminghaus T, Skarstad K. 2011. Replication patterns and organization of replication forks in *Vibrio cholerae*. Microbiology 157:695–708.

Sung W, Ackerman MS, Dillon MM, Platt TG, Fuqua C, Cooper VS, Lynch M. 2016. Evolution of the insertion-deletion mutation rate across the tree of life. G3-Genes Genomes Genet. In Press.

Sung W, Ackerman MS, Gout J-F, Miller SF, Williams E, Foster PL, Lynch M. 2015. Asymmetric context-dependent mutation patterns revealed through mutation-accumulation experiments. Mol. Biol. Evol. 32:1672–1683.

Sung W, Ackerman MS, Miller SF, Doak TG, Lynch M. 2012. Drift-barrier hypothesis and mutation-rate evolution. Proc. Natl. Acad. Sci. U. S. A. 109:18488–18492.

Sung W, Tucker AE, Doak TG, Choi E, Thomas WK, Lynch M. 2012. Extraordinary genome stability in the ciliate *Paramecium tetraurelia*. Proc. Natl. Acad. Sci. 109:19339–19344.

Thompson FL, Iida T, Swings J. 2004. Biodiversity of *Vibrios*. Microbiol. Mol. Biol. Rev. 68.

Val ME, Skovgaard O, Ducos-Galand M, Bland MJ, Mazel D. 2012. Genome engineering in *Vibrio cholerae:* A feasible approach to address biological issues. Plos Genet. 8:e1002472.

Vos M, Didelot X. 2009. A comparison of homologous recombination rates in bacteria and archaea. ISME J. 3:199–208.

Waldor MK, RayChaudhuri D. 2000. Treasure trove for cholera research. Nature 406:469–470.

Wollenberg MS, Ruby EG. 2012. Phylogeny and fitness of *Vibrio fischeri* from the light organs of *Euprymna scolopes* in two Oahu, Hawaii populations. ISME J. 6:352–362.

Xu X, Peng M, Fang Z. 2000. The direction of microsatellite mutations is dependent upon allele length. Nat. Genet. 24:396–399.

Ye K, Schulz MH, Long Q, Apweiler R, Ning ZM. 2009. Pindel: a pattern growth approach to detect break points of large deletions and medium sized insertions from paired-end short reads. Bioinformatics 25:2865–2871.

Yildiz FH, Visick KL. 2009. Vibrio biofilms: so much the same yet so different. Trends Microbiol. [Internet] 17:109–118. Available from: <Go to ISI>://000264681800004

Zhang XL, Mathews CK. 1995. Natural DNA precursor pool asymmetry and base sequence context as determinants of replication fidelity. J. Biol. Chem. 270:8401–8404.

Zhu YO, Siegal ML, Hall DW, Petrov DA. 2014. Precise estimates of mutation rate and spectrum in yeast. Proc. Natl. Acad. Sci. U. S. A. 111:E2310–E2318.

